# Evolutionary Divergence and Structural Differentiation of Multiple Immunoglobulin M Genes in Gekkota (Squamata: Reptilia)

**DOI:** 10.64898/2026.02.13.705754

**Authors:** Francisco Gambón Deza

**Affiliations:** Independent research

**Keywords:** Immunoglobulin M, Gekkota, gene duplication, molecular evolution, antibody diversification, reptile immunology

## Abstract

Immunoglobulin M (IgM) is the most ancient and conserved antibody class in jawed vertebrates and is typically encoded by a single gene. In contrast, geckos and related lizards (infraorder Gekkota) possess multiple IgM genes within the immunoglobulin heavy chain locus. Here, we analyze 52 IgM constant-region sequences from 13 Gekkota species to clarify the evolutionary origin and functional consequences of this expansion. Phylogenetic reconstruction showed that IgM1 (the canonical form) is nearly monophyletic (86.7% clade purity), whereas internal-locus IgM2–6 variants display complex, lineage-specific duplication patterns. We identified 53 diagnostic amino acid positions distinguishing IgM1 from other variants, concentrated in CH1 (19 positions) and CH2 (25 positions). These differences are accompanied by a pronounced physicochemical shift in CH2: IgM1 carries a net positive charge (+2.01) while other IgMs are negatively charged (−2.13), a Δ of +4.14 charge units. Conservation analyses indicate stronger constraint on IgM1 in CH1/CH2, while internal-locus IgMs are more conserved in CH4, consistent with maintained polymerization function. Three-dimensional structural comparison of IgM1 and IgM4 supports functional divergence in assembly: IgM4 adopts an “open mouth” CH1–CH2 conformation with increased heavy–light chain contacts and a more electrostatically enriched interface, suggesting compensatory stabilization mechanisms. Together, these results support specialization of internal-locus IgMs through combined sequence and structural divergence.

## 1. Introduction

Immunoglobulin M (IgM) represents the most evolutionarily ancient antibody isotype, present in all jawed vertebrates from cartilaginous fish to mammals [6]. As the first antibody produced during a primary immune response and the predominant immunoglobulin in early vertebrate lineages, IgM plays fundamental roles in complement activation, pathogen neutralization, and immune regulation [2].

The typical vertebrate immunoglobulin heavy chain locus contains a single functional IgM gene, consisting of four constant domains (CH1-CH4) that mediate effector functions including polymerization into pentameric structures and interaction with the J chain [3]. However, recent genomic studies have revealed unexpected complexity in the immunoglobulin loci of certain reptilian lineages, particularly within the infraorder Gekkota (geckos and related lizards).

Gekkota represents a diverse clade of squamate reptiles comprising over 1,800 species distributed across tropical and subtropical regions worldwide [7]. Comparative studies of immunoglobulin loci in amphibians and reptiles have highlighted unexpected diversity in heavy chain organization and constant-region gene content across lineages [9, 8, 16]. In several gecko species, multiple IgM-like genes have been identified within the heavy chain locus, with one gene (designated IgM1) occupying the canonical position and additional genes (IgM2–6) located internally within the locus. The evolutionary origin, structural characteristics, and potential functional significance of these additional IgM genes remain poorly understood.

Gene duplication is a major driver of evolutionary innovation, providing raw material for functional diversification [15]. In the context of immunoglobulins, gene duplication has given rise to the diverse antibody isotypes observed in mammals (IgM, IgD, IgG, IgA, IgE) and has contributed to the expansion of variable region gene families [5]. Understanding whether the multiple IgM genes in Gekkota represent functional diversification or pseudogenization has important implications for our understanding of reptilian immunity and antibody evolution.

In this study, we performed a comprehensive comparative analysis of IgM sequences from 13 Gekkota species to address the following questions: (1) What is the evolutionary relationship between IgM1 and the internal locus IgMs? (2) Are there systematic structural differences between these groups that might indicate functional divergence? (3) Which protein domains show the greatest differentiation, and what might this suggest about altered functions?

## 2. Materials and Methods

### 2.1. Sequence Data

IgM constant region sequences were obtained from 13 species representing major Gekkota lineages: *Asaccus caudivolvulus*, *Correlophus ciliatus*, *Eublepharis macularius*, *Euleptes europaea*, *Gekko hokouensis*, *Gekko kuhli*, *Heteronotia binoei*, *Lepidodactylus listeri*, *Lepidodactylus lugubris*, *Mediodactylus kotschyi*, *Paroedura picta*, *Sphaerodactylus townsendi*, and *Teratoscincus roborowskii*. A total of 52 sequences were analyzed, comprising 13 IgM1, 13 IgM2, 11 IgM3, 9 IgM4, 5 IgM5, and 1 IgM6 sequences. The complete set of analyzed sequences is provided as Supplementary Data S1 (FASTA).

### 2.2. Multiple Sequence Alignment

Amino acid sequences were aligned using MAFFT v7.490 [11] with the L-INS-i algorithm optimized for sequences with conserved domains. The resulting alignment of 540 positions was filtered to remove columns containing more than 80% gaps, yielding a final alignment of 439 positions for downstream analyses.

### 2.3. Phylogenetic Analysis

Phylogenetic relationships were inferred using the Neighbor-Joining method [17] with distances calculated from the BLOSUM62 substitution matrix [10]. Tree construction and manipulation were performed using Biopython v1.81 [4]. Monophyly of IgM groups was assessed by identifying the most recent common ancestor (MRCA) of each group and calculating the proportion of group members within the MRCA clade (clade purity).

### 2.4. Conservation Analysis

Positional conservation was quantified using Shannon entropy [18]:

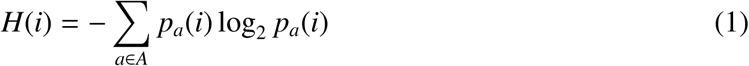

where *H*(*i*) is the entropy at position *i*, *A* is the set of amino acids, and *p_a_*(*i*) is the frequency of amino acid *a* at position *i*. Lower entropy values indicate higher conservation. Entropy was calculated separately for IgM1 sequences and for all other IgM sequences to identify differentially conserved regions.

### 2.5. Identification of Diagnostic Positions

Diagnostic positions were defined as alignment columns where: (1) the most frequent amino acid differed between IgM1 and other IgMs, (2) the dominant amino acid frequency exceeded 60% in IgM1 and 50% in other IgMs, and (3) the IgM1-characteristic amino acid occurred at less than 30% frequency in other IgMs and vice versa.

### 2.6. Physicochemical Property Analysis

Amino acid properties were characterized using the Kyte-Doolittle hydrophobicity scale [12] and standard charge assignments at pH 7.0 (K, R: +1; H: +0.1; D, E: −1). Mean hydrophobicity and net charge were calculated for each sequence and for each structural domain (CH1: positions 1-110; CH2: positions 111-220; CH3: positions 221-330; CH4: positions 331-439).

### 2.7. Three-Dimensional Structural Modeling and Interface Analysis

Representative IgM1 and IgM4 heavy-chain constant regions from *Eublepharis macularius* were modeled using AlphaFold2. Predicted heavy-chain models were combined with the corresponding lambda light chain to generate representative Fab-like assemblies, and structural comparisons were performed between IgM1 and IgM4. Interface residues and atomic contacts at the heavy–light chain interface were quantified from the modeled complexes, and geometric parameters were extracted, including the CH1–Light–CH2 angle, the distance between the CH2 domain center and the light chain center, and the number of CH2 residues participating in the interface. Interface composition was summarized by physicochemical class (hydrophobic, polar, positively charged, negatively charged) to evaluate the contribution of electrostatic interactions.

### 2.8. Three-Dimensional Structural Modeling and Interface Analysis

Representative IgM1 and IgM4 heavy-chain constant regions from *Eublepharis macularius* were modeled using AlphaFold2. Predicted heavy-chain models were combined with the corresponding lambda light chain to generate representative Fab-like assemblies, and structural comparisons were performed between IgM1 and IgM4. Interface residues and atomic contacts at the heavy–light chain interface were quantified from the modeled complexes, and geometric parameters were extracted, including the CH1–Light–CH2 angle, the distance between the CH2 domain center and the light chain center, and the number of CH2 residues participating in the interface. Interface composition was summarized by physicochemical class (hydrophobic, polar, positively charged, negatively charged) to evaluate the contribution of electrostatic interactions.

### 2.9. AI-Assisted Workflow

Parts of the computational workflow (including data handling, exploratory analyses, and code drafting) were assisted by guided AI agents implemented in the web platform Phylo.bio. All final analytical choices, parameter settings, and interpretations were reviewed and validated by the author.

### 2.10. AI-Assisted Workflow

Parts of the computational workflow (including data handling, exploratory analyses, and code drafting) were assisted by guided AI agents implemented in the web platform Phylo.bio. All final analytical choices, parameter settings, and interpretations were reviewed and validated by the author.

## 3. Results

### 3.1. Phylogenetic Relationships Among Gekkota IgM Genes

Phylogenetic analysis of 52 IgM sequences revealed distinct evolutionary patterns between IgM1 and the internal locus variants (Figure 1). IgM1 sequences formed a nearly monophyletic clade with 86.7% purity, meaning that 13 of the 15 sequences within the IgM1 MRCA clade were IgM1 (the two exceptions being IgM2 and IgM6 from *Eublepharis macularius*).

**Figure 1:**
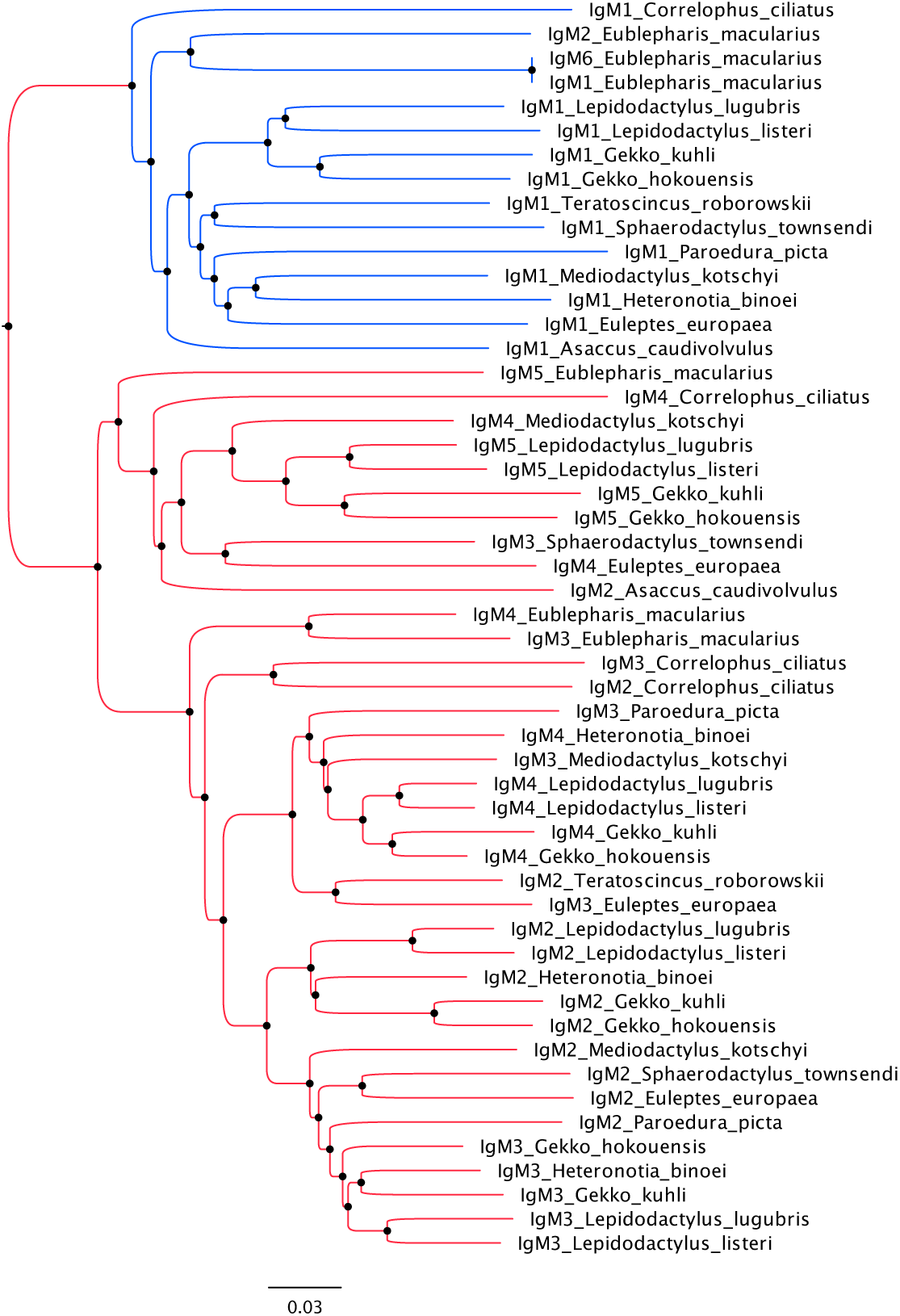
Phylogenetic tree of 52 IgM sequences from 13 Gekkota species. Sequences are colored by IgM type: IgM1 (yellow), IgM2 (blue), IgM3 (green), IgM4 (orange), IgM5 (pink), IgM6 (black). Note the near-monophyletic clustering of IgM1 sequences and the polyphyletic distribution of other IgM types.

In contrast, IgM2-6 sequences showed highly polyphyletic distributions. The MRCA of IgM2 sequences contained representatives of all IgM types (purity: 27.7%), and similar patterns were observed for IgM3 (21.2%), IgM4 (23.7%), and IgM5 (50.0%). This pattern suggests that while IgM1 diverged early and has been maintained as a distinct lineage, the internal locus IgMs arose through multiple independent duplication events specific to different Gekkota lineages.

### 3.2. Differential Conservation Patterns Across Domains

Conservation analysis revealed striking differences in evolutionary constraints between IgM1 and other IgMs across the four constant domains (Figure 2, Table 1).

**Figure 2:**
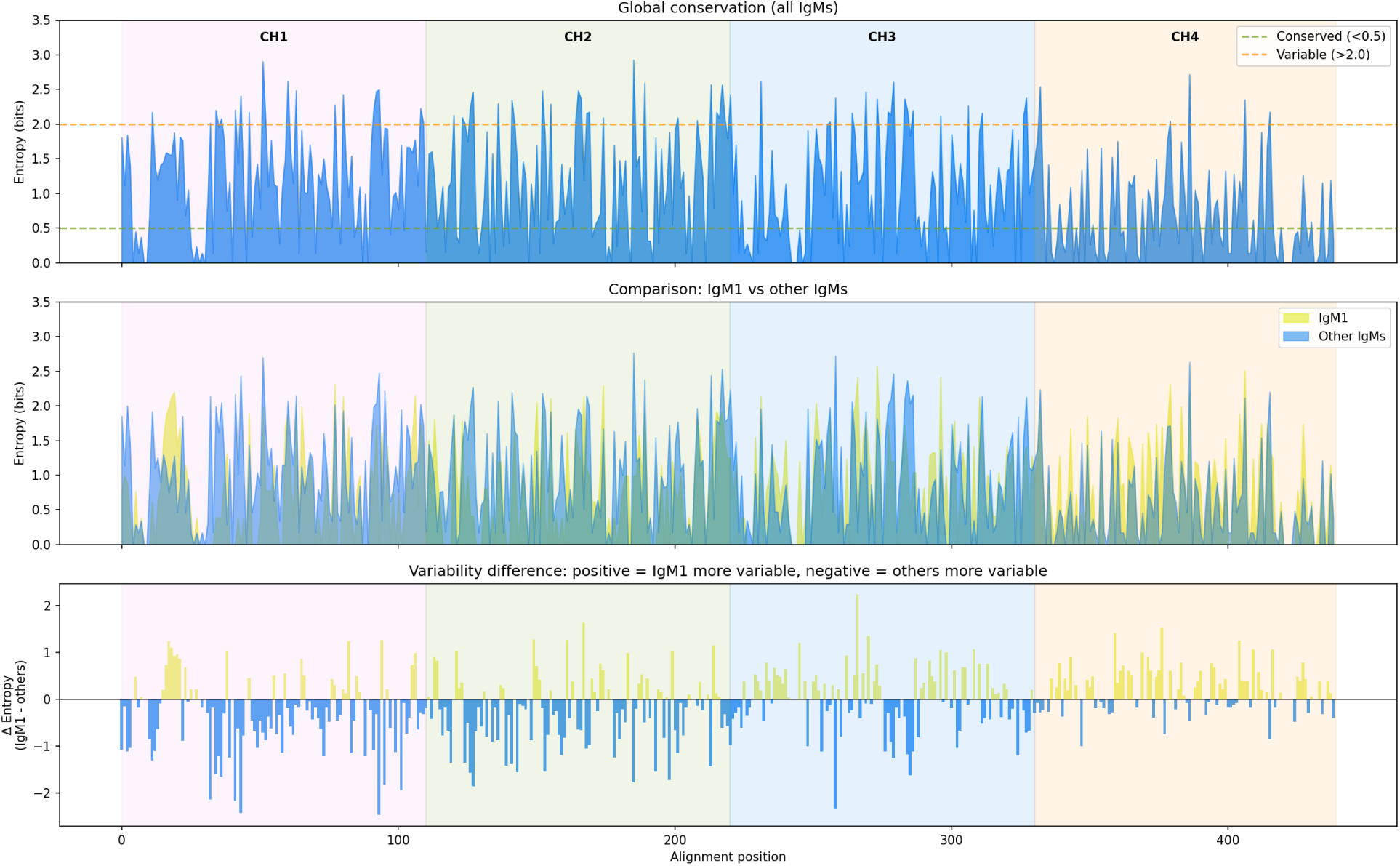
Conservation profile across the IgM constant region. (A) Shannon entropy for all sequences combined. (B) Comparison of entropy between IgM1 and other IgMs. (C) Difference in entropy (IgM1 minus others); positive values indicate positions where IgM1 is more variable. Approximate domain boundaries are indicated by colored shading.

**Table 1:** Mean Shannon entropy by domain and IgM group. Lower values indicate higher conservation.

| Domain | IgM1 | Other IgMs | $\Delta$ | More conserved |
| --- | --- | --- | --- | --- |
| CH1 (1-110) | 0.72 | 1.03 | -0.31 | IgM1 |
| CH2 (111-220) | 0.75 | 1.00 | -0.25 | IgM1 |
| CH3 (221-330) | 0.88 | 0.88 | 0.00 | Similar |
| CH4 (331-439) | 0.74 | 0.57 | +0.17 | Other IgMs |

IgM1 sequences showed significantly lower entropy (higher conservation) in the CH1 and CH2 domains compared to other IgMs. In CH1, mean entropy was 0.72 bits for IgM1 versus 1.03 bits for other IgMs. In CH2, values were 0.75 versus 1.00 bits, respectively. This pattern was reversed in CH4, where other IgMs showed higher conservation (0.57 bits) compared to IgM1 (0.74 bits). The CH3 domain showed similar conservation levels in both groups (0.88 bits).

### 3.3. Diagnostic Amino Acid Positions

We identified 53 diagnostic positions that systematically distinguish IgM1 from other IgM variants (Table 2). These positions were unevenly distributed across domains: CH1 contained 19 diagnostic positions (35.8%), CH2 contained 25 (47.2%), CH3 contained 6 (11.3%), and CH4 contained only 3 (5.7%).

**Figure 3:**
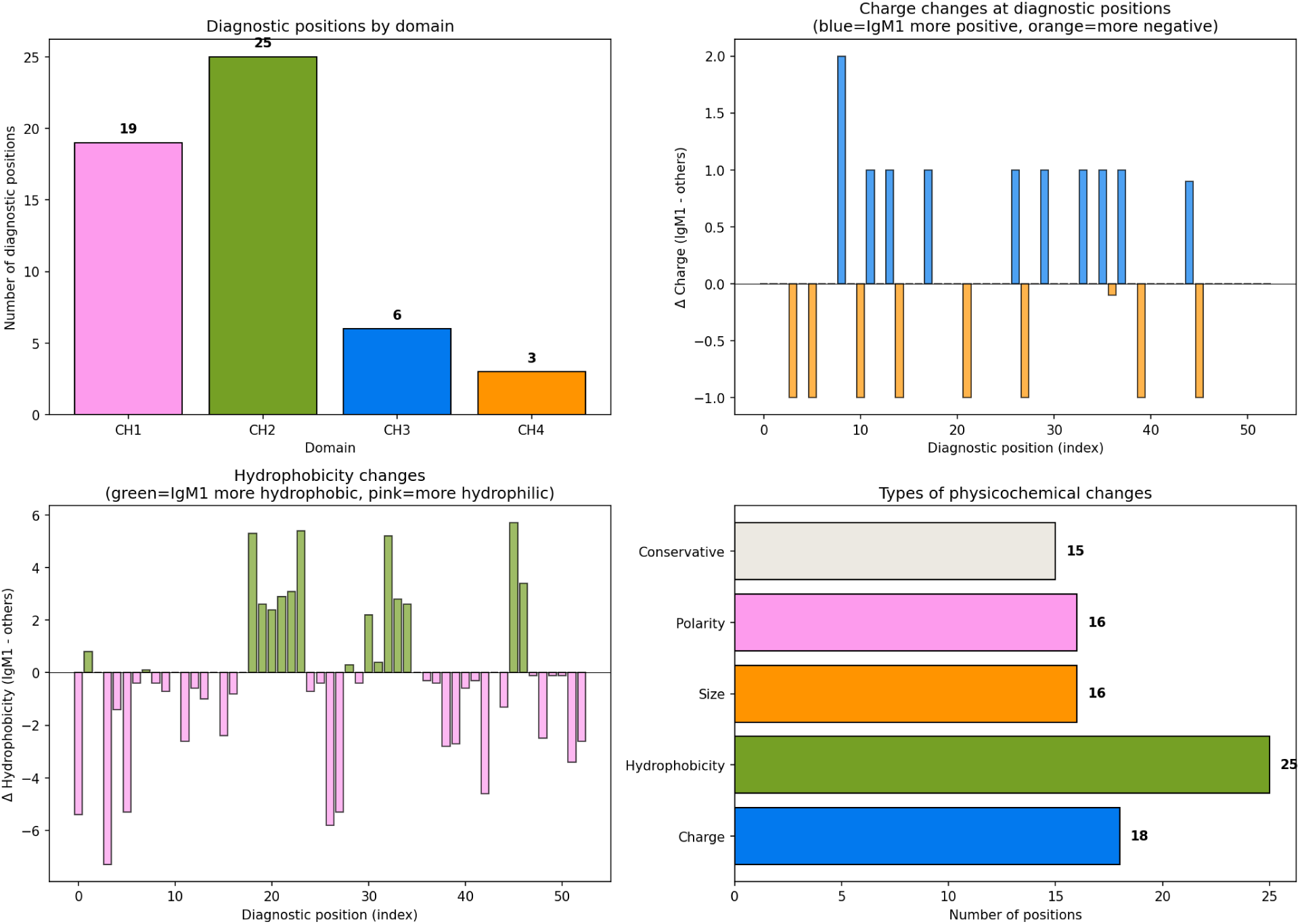
Analysis of diagnostic positions. (A) Distribution of diagnostic positions by domain. (B) Charge changes at diagnostic positions. (C) Hydrophobicity changes at diagnostic positions. (D) Summary of physicochemical change types.

**Table 2:** Distribution of diagnostic positions and types of physicochemical changes by domain.

| Domain | Diagnostic positions | Charge changes | Hydrophobicity changes | Size changes | Polarity changes |
| --- | --- | --- | --- | --- | --- |
| CH1 | 19 | 8 | 6 | 4 | 5 |
| CH2 | 25 | 8 | 14 | 6 | 6 |
| CH3 | 6 | 2 | 3 | 1 | 1 |
| CH4 | 3 | 0 | 2 | 1 | 0 |
| **Total** | **53** | **18** | **25** | **12** | **12** |

Among the 53 diagnostic positions, 18 involved changes in amino acid charge, 25 involved hydrophobicity changes, 12 involved significant size differences, and 12 involved polarity changes. Notably, 15 positions showed conservative substitutions with minimal physicochemical impact.

### 3.4. Critical Charge Inversion in the CH2 Domain

The most striking finding was a dramatic difference in net charge of the CH2 domain between IgM groups (Figure 4). IgM1 sequences exhibited a mean net charge of +2.01 in CH2, while other IgMs showed a mean charge of −2.13, representing a difference of +4.14 charge units (Table 3).

**Figure 4:**
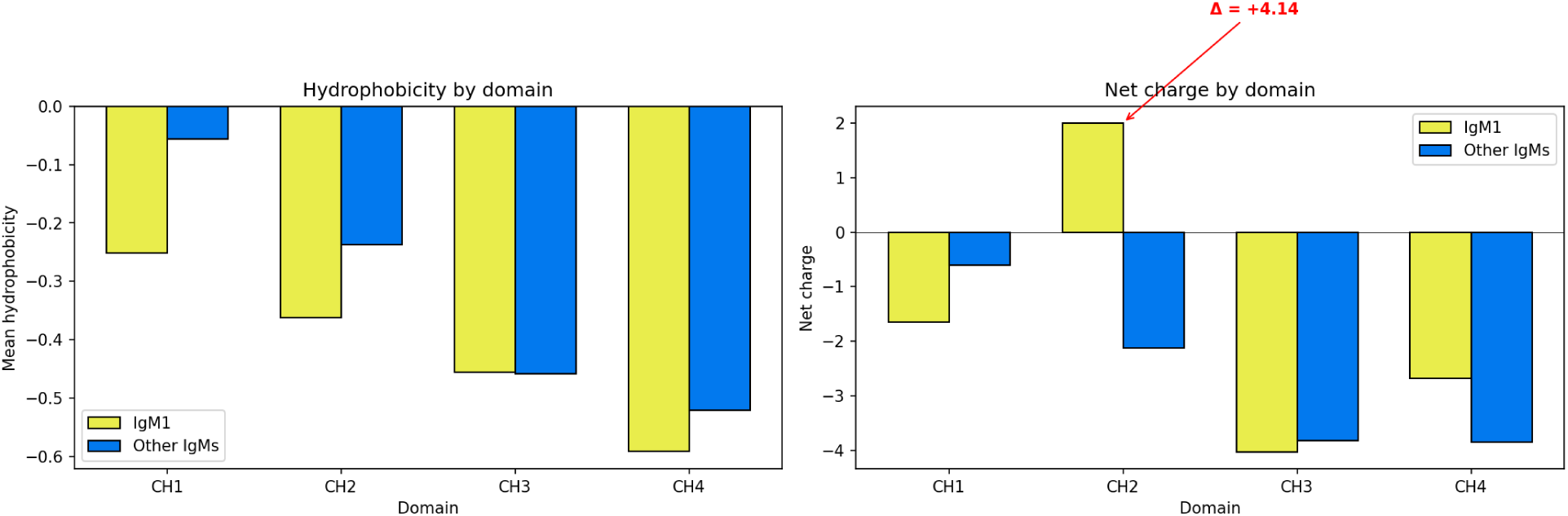
Physicochemical properties by domain. (A) Mean hydrophobicity comparing IgM1 and other IgMs. (B) Net charge by domain, highlighting the dramatic charge inversion in CH2.

**Table 3:**
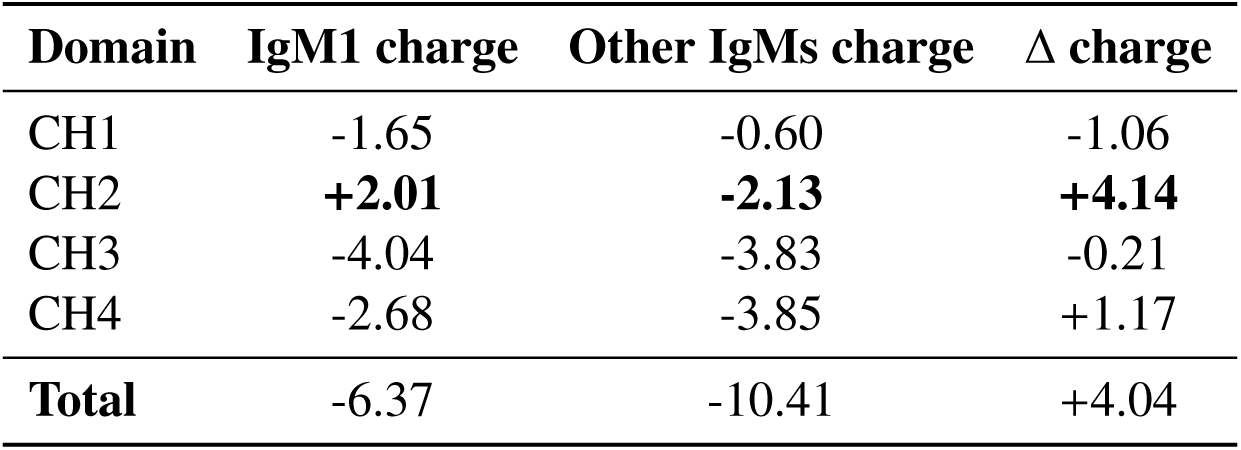
Net charge by domain comparing IgM1 and other IgM variants.

| Domain | IgM1 charge | Other IgMs charge | $\Delta$ charge |
| --- | --- | --- | --- |
| CH1 | -1.65 | -0.60 | -1.06 |
| CH2 | **+2.01** | **-2.13** | **+4.14** |
| CH3 | -4.04 | -3.83 | -0.21 |
| CH4 | -2.68 | -3.85 | +1.17 |
| **Total** | **-6.37** | **-10.41** | **+4.04** |

This charge inversion results from the cumulative effect of 8 diagnostic positions in CH2 involving charged amino acids. At position 119, IgM1 has proline while other IgMs have arginine. At positions 135, 147, 181, 194, and 199, IgM1 has positively charged or neutral residues where other IgMs have neutral or negatively charged residues.

### 3.5. Global Sequence Properties

Comparison of overall sequence properties revealed that IgM1 sequences are slightly shorter (413.2 ± 20.4 aa) than other IgMs (428.0 ± 15.2 aa), more hydrophilic (mean hydrophobicity: −0.41 vs −0.32), and carry less negative net charge (−6.37 vs −10.41). The proportion of positively charged residues was higher in IgM1 (11.28%) compared to other IgMs (10.43%).

### 3.6. Three-Dimensional Structural Analysis

To validate the sequence-based predictions and understand the structural basis of the observed differences, we analyzed the predicted three-dimensional structures of IgM1 and IgM4 from *Eublepharis macularius* as homodimers with lambda light chains (Figure 5). Additional three-dimensional models for IgM1, IgM4 and IgM5 are provided as Supplementary Figure S1 (Figure 6).

**Figure 5:**
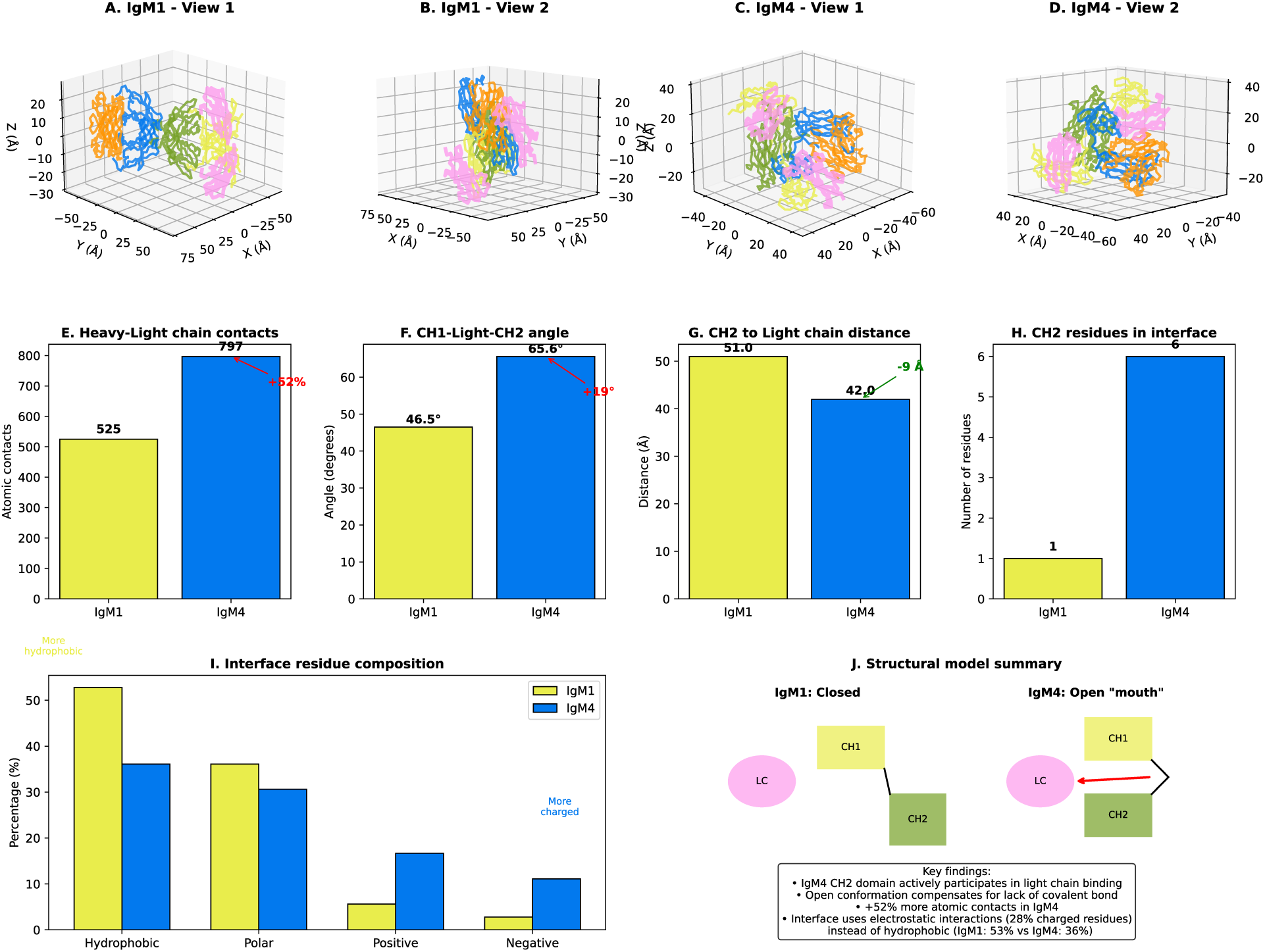
Three-dimensional structural comparison of IgM1 and IgM4 from *Eublepharis macularius*. (A-B) IgM1 structure shown from two orientations, displaying the closed conformation where CH2 (green) is distant from the light chain (pink). (C-D) IgM4 structure showing the open “mouth” conformation where CH1 (yellow) and CH2 (green) together embrace the light chain. (E) Comparison of atomic contacts at the heavy-light chain interface. (F) CH1-Light-CH2 angle comparison showing the wider angle in IgM4. (G) Distance from CH2 domain center to light chain center. (H) Number of CH2 residues participating in the interface. (I) Composition of interface residues by physicochemical class. (J) Schematic model summarizing the structural differences. Domain colors: CH1 (yellow), CH2 (green), CH3 (blue), CH4 (orange), Light chain (pink).

#### 3.6.1. Differential Heavy-Light Chain Interface

Structural comparison revealed striking differences in how IgM1 and IgM4 interact with the light chain (Table 4). Despite having similar numbers of interface residues (36 in both cases), IgM4 exhibited 52% more atomic contacts (797 vs 525), indicating a more extensive interaction surface.

**Table 4:** Structural comparison of heavy chain-light chain interface between IgM1 and IgM4.

| Parameter | IgM1 | IgM4 | Difference |
| --- | --- | --- | --- |
| Interface residues | 36 | 36 | – |
| Atomic contacts | 525 | 797 | +52% |
| CH1-Light-CH2 angle | 46.5° | 65.6° | +19° |
| CH2-Light distance | 51.0 Å | 42.0 Å | -9 Å |
| CH2 residues in interface | 1 | 6 | +5 |
| *Interface composition (%)* |  |  |  |
| Hydrophobic | 52.8 | 36.1 | -16.7 |
| Polar | 36.1 | 30.6 | -5.5 |
| Positively charged | 5.6 | 16.7 | +11.1 |
| Negatively charged | 2.8 | 11.1 | +8.3 |
| Total charged | 8.4 | 27.8 | +19.4 |

#### 3.6.2. The “Open Mouth” Conformation of IgM4

The most notable structural difference was the geometry of the CH1-CH2 domains relative to the light chain. In IgM1, the CH2 domain is positioned distally from the light chain (51.0 Å), with only one CH2 residue participating in the interface. The angle formed by CH1-Light chain-CH2 is relatively acute (46.5°), representing a “closed” conformation typical of classical immunoglobulins. In contrast, IgM4 adopts an “open mouth” conformation where both CH1 and CH2 domains actively participate in light chain binding. The CH2 domain is positioned 9 Å closer to the light chain (42.0 Å), and six CH2 residues (positions 137, 194-198) directly contact the light chain. The CH1-Light-CH2 angle is significantly wider (65.6°), creating a cleft that embraces the light chain from two sides.

#### 3.6.3. Electrostatic Compensation for Non-Covalent Association

The interface composition analysis revealed a fundamental shift in the nature of heavy-light chain interactions. IgM1 relies predominantly on hydrophobic contacts (52.8% of interface residues), consistent with the classical immunoglobulin fold where a conserved disulfide bond provides covalent stabilization.

IgM4, lacking this covalent bond, has evolved an alternative strategy: the interface contains significantly more charged residues (27.8% vs 8.4%), suggesting that electrostatic interactions compensate for the absence of the disulfide bond. This finding directly explains the charge inversion observed in the sequence analysis, where IgM1 CH2 carries a net positive charge (+2.01) while IgM4 CH2 is negatively charged (−2.13).

#### 3.6.4. Structural Validation of Sequence Predictions

The structural analysis validates and extends our sequence-based findings:

1. The concentration of diagnostic positions in CH1/CH2 (83%) corresponds to the domains showing the greatest structural divergence in light chain interaction mode.
2. The higher hydrophobicity of IgM4 CH1/CH2 in sequence analysis reflects residues oriented toward the protein interior, not the interface, explaining why the interface paradoxically contains fewer hydrophobic residues.
3. The charge inversion in CH2 (Δ = +4.14) has clear structural significance: IgM4 uses electrostatic interactions at the extended CH2 interface to maintain light chain association.
4. The greater conservation of CH4 in internal locus IgMs may relate to maintained polymerization function, as this domain shows no involvement in the altered light chain binding mode.

## 4. Discussion

### 4.1. Evolutionary Origin of Multiple IgM Genes in Gekkota

Our phylogenetic analysis provides strong evidence for a two-phase model of IgM diversification in Gekkota. The near-monophyly of IgM1 (86.7% clade purity) indicates that this gene represents the ancestral IgM that has been maintained under purifying selection across all Gekkota lineages. The high conservation of IgM1, particularly in the CH1 and CH2 domains involved in antigen binding and effector function initiation, suggests that IgM1 retains the classical immune functions of vertebrate IgM.

In contrast, the polyphyletic distribution of IgM2-6 sequences indicates that these genes arose through multiple independent duplication events in different Gekkota lineages rather than from a single ancestral duplication. This pattern is consistent with ongoing “birth-and-death” evolution [13] of immunoglobulin genes, where gene duplications occur repeatedly and duplicated copies either acquire new functions, become pseudogenes, or are lost.

### 4.2. Functional Implications of Structural Divergence

The concentration of diagnostic positions in CH1 and CH2 (83% of all diagnostic positions) suggests that these domains have experienced the strongest divergent selection between IgM1 and other IgMs. The CH1 domain mediates interaction with the immunoglobulin light chain, and changes in this region could affect antibody assembly or stability. The CH2 domain in IgM corresponds to the hinge region in other immunoglobulin classes and influences molecular flexibility and the spatial arrangement of antigen-binding sites.

The dramatic charge inversion in CH2 (+2.01 in IgM1 vs −2.13 in other IgMs) is particularly intriguing. Electrostatic properties of antibody domains influence protein-protein interactions, including binding to Fc receptors and complement components [14]. A positively charged CH2 in IgM1 might favor interactions with negatively charged cell surface molecules or complement proteins, while the negatively charged CH2 in other IgMs could alter or abolish such interactions.

### 4.3. Potential Specialized Functions of Internal Locus IgMs

The higher conservation of CH4 in internal locus IgMs compared to IgM1 suggests that polymerization function may be particularly important for these variants. The CH4 domain mediates IgM polymerization through interactions with the J chain and between IgM monomers [3]. Enhanced conservation of CH4 could indicate selection for efficient polymerization, possibly related to specialized functions in mucosal immunity where polymeric immunoglobulins play important roles.

Several hypotheses could explain the maintenance of multiple IgM genes in Gekkota:

1. **Compensatory assembly mechanisms**: The “open mouth” CH1–CH2 conformation in IgM4, together with increased atomic contacts and an enrichment of charged residues at the heavy–light chain interface (Figure 5), suggests selection for alternative stabilization strategies when the canonical inter-chain disulfide bond is absent or weakened. This points to diversification in antibody assembly constraints rather than simple redundancy.
2. **Modulation of Fab geometry and binding dynamics**: The wider CH1–Light–CH2 angle observed in IgM4 implies altered Fab geometry and potentially different conformational flexibility. Such changes could affect antigen-binding kinetics and accessibility to epitopes, providing a mechanism for functional divergence without necessarily altering polymerization.
3. **Electrostatic re-tuning of e**ff**ector interactions**: The marked CH2 charge inversion between IgM1 and other IgMs may impact interactions with complement components and/or Fc receptor-like molecules. The structural data support an adaptive role for charge redistribution at exposed surfaces and interfaces, consistent with altered downstream effector engagement.
4. **Subfunctionalization with preserved polymerization**: Strong conservation of CH4 among internal locus IgMs is consistent with maintaining polymerization-related functions, while divergence concentrated in CH1/CH2 supports partitioning of functions tied to light-chain association and Fc-proximal properties between IgM1 and internal locus IgMs.
5. **Regulatory specialization (expression context)**: Differential expression across tissues, developmental stages, or immune contexts may still contribute to maintaining multiple copies, but the structural divergence indicates that expression differences likely act on proteins with distinct biophysical properties rather than near-identical isoforms.

### 4.4. Comparison with Other Vertebrates

Multiple IgM genes have been reported in some teleost fish species, where they appear to have arisen through lineage-specific duplications similar to what we observe in Gekkota [1]. However, the pattern in Gekkota is unusual among amniotes, where typically only a single functional IgM gene is present. The independent evolution of multiple IgM genes in fish and geckos suggests that this represents a convergent evolutionary strategy, possibly related to similar ecological or immunological pressures.

### 4.5. Structural Basis for Functional Divergence

The three-dimensional structural analysis provides a mechanistic explanation for the sequence divergence patterns observed between IgM1 and internal locus IgMs. The “open mouth” conformation of IgM4, where CH2 actively participates in light chain binding, represents a novel structural adaptation not previously described in immunoglobulins.

This conformational difference has several functional implications:

#### Light chain association stability

The 52% increase in atomic contacts and the involvement of CH2 in the interface suggest that IgM4 may have comparable or even enhanced light chain binding stability despite lacking the canonical inter-chain disulfide bond. The use of electrostatic interactions (27.8% charged residues vs 8.4% in IgM1) provides an alternative stabilization mechanism.

#### Conformational flexibility

The wider CH1-Light-CH2 angle in IgM4 (65.6° vs 46.5°) may confer different dynamic properties to the Fab region, potentially affecting antigen binding kinetics or the range of accessible epitopes.

#### Evolutionary trajectory

The structural data suggest that following gene duplication, internal locus IgMs underwent substantial conformational evolution in the CH1-CH2 region while maintaining the CH3-CH4 architecture required for polymerization. This is consistent with sub-functionalization, where the ancestral IgM functions have been partitioned between IgM1 (classical antigen binding) and internal locus IgMs (potentially specialized for different immune contexts).

The discovery of this “open mouth” conformation in reptilian IgMs adds to the growing appreciation of structural diversity in vertebrate immunoglobulins and suggests that alternative solutions to antibody architecture may be more common than previously recognized.

### 4.6. Limitations and Future Directions

This study focused on sequence-based analysis and did not include experimental validation of the proposed functional differences. Future studies should investigate:

- Expression patterns of different IgM genes across tissues and developmental stages
- Biochemical characterization of polymerization efficiency and complement activation
- Structural determination of representative IgM variants to map diagnostic positions in three-dimensional context
- Population-level variation to assess ongoing selection pressures

## 5. Conclusions

Our comprehensive analysis of IgM diversity in Gekkota reveals a complex evolutionary history involving an ancestral IgM1 that has been maintained under strong purifying selection, alongside multiple lineage-specific duplications that have given rise to structurally divergent IgM variants. The concentration of diagnostic changes in CH1/CH2 domains and the dramatic charge inversion in CH2 suggest functional divergence between IgM1 and internal locus IgMs. These findings provide a foundation for understanding the unique immunoglobulin repertoire of geckos and highlight the ongoing evolutionary plasticity of the vertebrate immune system.

## Supporting information

latex

## Supplementary Material

### Data S1. IgM sequences (FASTA)

#### >IgM1_Asaccus_caudivolvulus

SQTAPSLFPLIPSWDTDTTVNGDVTVGCMAKDFLPDSLTLSWQKPNNQSLEANRIRRFPS FAKSNGLNTACSQATVPADQWKAFEPFYCKAELASQNTVTRVVRQIVCLPPEIVIRVPPL EDFHGAYLNATLLCKADNLHTKQTTIKWLEDGNVLDSGFTTSAPIQQGNGYSIVSELIVT KKDWYTEKTYSCQVKNEKFNEIRNVSKSSVCEGCDGTNVPVRVETIPPSFEDIFLTKSAI LTCRISNIPYSQDIAELKVSWLREYNNTRLETKIGQATGQENKDLWFVDATTTICAEEWQ KGETYLCQVQLPLLASTETKSLKKATGSTYLPPAVYVLPPPSEQLALQETATITCLLKGF YPNNFFVKWLRNNEQVAASEYFTSQPVQESKSPERYFTYSTLNVKEQDWSYGATYTCVVG HEALPLQTTQKSVDKNT

#### >IgM2_Asaccus_caudivolvulus

TPSGPFLFPLLLPAENTPSNGLVTVGCLAKDVPPALVRFTWENQNNVSINASYVKQYPAL SISPGTFTASSLVAVPADAWENSEAFYCKAESVDGAARVVRQASPTEPNMIIRAPRPKDF DNSDGGATIFCKAEHLPTAGTTIQWLKDGNVVESGFTTTEPVGMRCSGYSIQSELVVDKD DWNADRKFSCRAHSNEFSETQKISKCMLCDGCSDTAAVHVETIPPSFADIYTTNSAKLTC RISNIPYGETLTKWNVTWTRESDSKELETVIGEAKEQENSGLLFVDVTTTICKEEWESQS TFACKVNLPRLLPTTETKTLRKPHGGTPKAPSVYVLPPPSEQLALQETATITCLLKDFYP NNFFVKWLRNNEPVGDSEYFTSQPIQESKSPERYFTYSTLNVREQDWSYGATYTCVVGHE ALPLQTTQKTIDKNT

#### >IgM1_Correlophus_ciliatus

SQSAPSLFPLIPSTSSIPDSGDITIGCLAKDFLPDTLTFSWQKPDNGSLEPSSVKRFPSM SSSGTFTVSSEATVPSGQWKTFEPFYCKAEHPSATKVARVVRQMTCEDPEVTVRVPPLEE FLGPYQNATILCKADHLHTERTTIKWLMNGKELTSDFTTGAPFKGRGGFSIISELIVTKR NWFAEKVFSCQVQNEKINLIRNVSKASVCQVRVETIPPSYADIFLNQSVKLTCRVSNLPY GPDGVDMLNVSWTRSPDNKPLETIVGEPKEQEGSALVFVEATATICLEEWNRGDTFTCKV THPLLVTTETKTLKKQIGGPRKAPSVYVLPPPSEQLVLHEKATLTCLLEGFFPGDFFVQW IWNNKPVDPSEYHTSKPIKGSKTTETYFAYSTLNVDEQEWSSGAQATCMVGHEAIHFQTT QKTIDKNT

#### >IgM2_Correlophus_ciliatus

PQGAPSLFPLIQTSERVSRDGLITVGCLAKDVVPQVTSFSWNDQNNVSVNANYIRQFTSV FNPNGTFTSSSQVVISASDFWDFVPFYCKATNTAGTSVARIARQTSCQDPEMIIRAPRVE DFESSDPNATIFCMAVNLHSESTTIQWLKNGQVIKSGFDTTGPVAAGRSGYSIMSELTVD KRDWVSDKKFSCEVHSETFTSIKNISKSLVCDLGPSVEVNVHVETIPPTFTDIYLTQSAK MTCRISNLPYDQNLTDLVVTWTRASDNKQLDTIQGSLQAQPNNEFGFVEATATVCPEEWN SGQTFNCKVTFPSVLPTAETKPLKKLKEGTPHAPAVYVLPPPSEQLILRETATVTCLIKG FYPNDFFVKWLRNDEPVLDSEYHTGAPLKESKTPETYFAYSTLNVNEQDWSYGAIYTCVV GHEELPLQTTQKTIDKNT

#### >IgM3_Correlophus_ciliatus

NQTAPSLFPLILPSESIPSNGLTTIGCLAKDVLPALVSFSWDDYSNVSINADHIMQYPTI RSSPDMYSSISRVAVSANDWRNFEPFYCKAEHVAGVARIVRQGKIIWFLLFDFERSDP NATIFCMATNLHTASTTLQWLKNGQALKSGFATTGPVAVGHSDYSIISELIVDRRDWVSD KTFSCEVHNENFSGMKNISKSLVCDLGPSTDVKLHVQTFPPTFTDIYLTKSAKLTCRISN IPYDNKLEDLEVIWTRASDNKQLDTIRGKLQAQQNTEYAFLDATASVCPEEWGSGQTFNC KVTFPSVLPTAETKPLRKLKEGTPSPPAVYVLPPPSEQLILRETATVTCLIKGFYPNDFF VKWLRNDEPIQPSQYHISNPIQESKTPERYFTYSTLNVNEQDWSYGAIYTCVVGHEELPL QTTQKTIDKNT

#### >IgM4_Correlophus_ciliatus

VPSLFPLILPTDSIASGEPITIGCLAKDALVRFTWTDYNNANISASDIKQYPTISTTAGT FIASTRVAVSANDWENFKPFYCTAEGVGKARLIRPKTNQPDVTIFCMANHLRTASATIQW LKEGNVLDAGFNTTGPGSIGCSGHSIFSKLTINKSDWDADKKFTCLVQNGEFRDLRNISK TSMCGKVKKSIPCCFDPQVKVETIPPSFVDIYLTQSANLTCRISNLPSGPDDLDMNVSWT RTSDNKELETVIGEAKQQENSGLFFVDATATICKEEWESGESFQCKVVADSLLPSAETRI LRKLHGGTPHAPSVYVLSPPSEQLALGETATMTCLLKNFYPKEFFVKWLRNDEPVDSENH TSDPIQESKTPERYFAYSTLNVSEQDWRHGATYTCVVGHEALPYQTTQKTIDKNM

#### >IgM1_Eublepharis_macularius

PQSAPFLIPLIPSSITDSEEIAVGCLAKDFLPNSLILSWQKPNNESLEAAKIRQFPSIFN SVGAFTASSEATIPARQWQAFEPFYCKAELASETRVARLVHQTPIACLPPEITIRVPPLD DFRGAFLNATILCKADHLHTRHTAIKWLKDGKVLTSGFTTTTSTRRGRDGYTIISELTVT KKDWFAEKIFSCQVQNEKINETRNVSKPSACEGCDCNVPIHVETIPPPFSDIYLTKSARL TCRISNIPYSQDLAELNVTWTRASDNKPLETVIGQAKEQENREMLFVDATVTLCPDEWEN GSTYKCKVTLPLMAKAEVKSLKKQNGGSRRAPSVYVLPPPPEQLALKGIATVTCLLKSFY PSDFFVKWLQNGEPVGVSEYYTSNPIQDSRTPKRYFSYSMLNVDEQDWSSGVTYTCVVGH EAFPVQTIQKTIDKDT

#### >IgM2_Eublepharis_macularius

SQSAPSLFPLIPSSSITDSNEITIGCMAKDFLPDSLTLSWQKPNNDSLEADKIKRFPSIN NPTGTFTASSAATVPTSQWDAFKPFYCKAEHASTTKVARVVRQTPIVPIEPTIIIRVPPL EDFLGPYLNASLLCKADHLYTEKTTISWMKNGQVVISGITTTAPIRQNSGFSIISELTVT KKDWYADTEFSCQVQNEKFNEIRNVSKASVCEGGGECNVRIRVETIPPSFNDIYLTKSAT LTCRISNIPFGEDLALNVTWTRASDKKTLETTTGQAKEQENRELVFVDATATICAEEWEK GDTYTCTVKHRLLATTEVKSLKKQNGRLFYIIPQNSGSHSAPAVYVLPPPSEQLALQET ATLTCLVKGFYPSDFFVKWLQDGEPVPVSEYFTSGPIQESRSPERYFAYSTINVNEQDWN SGDHYTCVVGHEALPLQMTQKTVDKHT

#### >IgM3_Eublepharis_macularius

SPSAPSLFPLILPSERLPSNGRITIGCLAKDVVPPVDSFTWDDLNNVSINANYIKQFPSV FNPNGTFTLSSQVSVSVNDWRNFEPFFCKANNTVGTTLARVTRQTSCLKPELNIHAPRLE DFENPDGNATLVCIAVNLHTPSTTVQWRKNGHVLDSGFTTTGPVAVGPSGYSIVSELTVD KRDWMTGKPYSCEVHNEEFSDIKNISKPLTCDSGPCSDAQVHVETIPPSFTDIYLTKSAK LTCRISNIPYDQDLKVLDVIWTRKSDNKELKTTKGEKQEQENNELVFVDATASVCVEEWD SGETFECKVTLPLLPTTEIKTLTKPKGGTPRAPEVYVFPPPSEQLKLQETATLTCLLKGF YPSDFFVTWLRNEEPVGDSEYITSHPIQESKTPERYFTYSTLNVKEQDWSYGATYTCVVG HEALPLQTIQKTIDKHT

#### >IgM4_Eublepharis_macularius

NQTSPSLFPLILPSERIPSNGLITIGCLAKDVLPALVSFSWDDSNNISLNTNYVTKYPTI GNPAGTFSSVSQVAVSANDWRNFQPFYCKAENVDGVARVVRHTSCIEPEMIVRAPRLEDF ENSDSNATLICIAVNLHTLSTTVQWRKNGHLLDSGFTTTGPVAVGHSGYSIVSELTVDRR DWVTEKKFSCEVHNEKFSGIKNISKSLVCDLGPCSDAQVHVETIPPSFTNIYLTKSAKLT CRISNMPYDQDLEVLDVIWTRKSDNKELKTTKGEKQEQENNELVFVDATASVCVEEWDSG ETFECKVTLPLLPAAEKKTLQKPKGGTPRAPEVYVLPPPSEQLKLQETATLTCLLKGFYP SDFFVTWLRNNEPVGASEYITSHPIQESKTPERYFTYSTLNVKEQDWSYGATYTCVVGHE ALPLQTIQKTIDKHT

#### >IgM5_Eublepharis_macularius

TQSGPSLFPLILPPSENTPCNGLVTIGCLAKDVPPALVRFTWDNHNNVSINASYVKQYPT IRNSAGTFTTSSQVVVSANDWQNFEPFYCKAEPVVGVARVVRHSKMASDMEPEMIIRAP CREDFENSNRTANIFCMAANLRTERTTLQWLKDGEILDSGFTTTGPVDVACSGFSIFSEL TITKNDWITDKKFSCRVHNKRFGDIRNISKSLVCDFGGDGNMPIHMETISPSFTDIYLTK AAKLTCRISNIPYSQDLATLNISWTRKRDNKPLETITGQAKQQENKDLVFVDATATVCTE EWESGDTFECKVTFSLLPAAEIRTLRKFQGGTPRAPAVYLLPPPSEQLALQETATLTCLL KGFYPNDVFVKWLHDGEPVGVSEYHTSSPIEESKSPELYFTYSTLTIDEQAWSSGTSYTC VVGHEALPLQTTQKTVDKNT

#### >IgM6_Eublepharis_macularius

PQSAPFLIPLIPSSITDSEEIAVGCLAKDFLPNSLILSWQKPNNESLEAAKIRQFPSIFN SVGAFTASSEATIPARQWQAFEPFYCKAELASETRVARLVHQTPIACLPPEITIRVPPLD DFRGAFLNATILCKADHLHTRHTAIKWLKDGKVLTSGFTTTTSTRRGRDGYTIISELTVT KKDWFAEKIFSCQVQNEKINETRNVSKPSACEGCDCNVPIHVETIPPPFSDIYLTKSARL TCRISNIPYSQDLAELNVTWTRASDNKPLETVIGQAKEQENREMLFVDATVTLCPDEWEN GSTYKCKVTLPLMAKAEVKSLKKQNGGSRRAPSVYVLPPPPEQLALKGIATVTCLLKSFY PSDFFVKWLQNGEPVGVSEYYTSNPIQDSRTPKRYFSYSMLNVDEQDWSSGVTYTCVVGH EAFPVQTIQKTIDKDT

#### >IgM1_Euleptes_europaea

SPSAPSLFPLIPSSEDTITGDSITVGCLAKDFLPDSLTLSWQKPNNESLEAAKISRFPSI MKTSGVFTASSEATVPADQWEAFEPYFCKAELNSQTKVARVVRQKDFRGAYLNATLFCKA DNLHTERTTMKWLEGGNVLNSGFTTTAPVRQGRNGYSMSSELIVTKKDWYADKIYSCQVQ NDKFNEIRNVSKPSVCEGCDGTTVPVHVETIPPPFFDIFHSKSAKLTCRISNLPLGQDLK EINVTWTREHDGKELEATFGQAEQQENRELVFVDVTANICAEEWESGDNYLCKVALPLLA STETKPLKKQNGGSHNPPSVYILPPPSEQLALQETATITCLLKGYFPNNFFVKWLRNDEQ VADSEYSVSTPIQESKSPERYFTYSTLNINEQEWTAGDRFTCVVGHEALPLQTTQKTVDK QT

#### >IgM2_Euleptes_europaea

PPSAPSLFPLILPSESIPNSGLITIGCLAKDVVPPVVSFSWDDQNVSINGSYIKQFPSVF NPMGTFTASSQVAISANDWWNFQPFYCRANNANGTGVAQVIRPTSCLEPEMIIRAPRLED FENSNPNASIVCMATNLHTASATVQWWKNGHVLGSGFTTTGPAVMGRGGYSIVSELEVDK RDWVSDKKFSCEVRSENFTSTKNISRPLVCDLGIVPDAKIHVATIPPSFTDIYLTKSATL TCRISNIPYDHDIKELDVVWTRERDNTELETRIGQATPQENRELVFVDATTTVCPDEWNS GETFKCKVAIPSLLPTAETRALKKLQGSTYRAPAVYVLPPPSEQLALKETATLTCLLKGY FPNDFFVKWMRNDEPVADSEYFTSKPIQESRTPELYFTYSTLNVREQDWSYGAIYTCVVG HEALPLQTTQKTIDKNT

#### >IgM3_Euleptes_europaea

TQTGPSLFPLVSRYESIPSNGLMTIACLAKDVPPAFVNFSWDDYNNVSINAKYVTRYPTI SNSAGTFSSVSQVAVSANDWRNFQPFYCKAEHVNGTARVVRQTSCIEPEVIIRAPRLQDF EAANPNATIVCMAVNLYTTSTTVQWLKNGHAVDSDFANTGPVAVGPHGYSIISELIVDRR DWVSDKTFSCEVHNENFTSIKNISKALVCDLGSCSDVQVHVETMPPSFTEIYLTKSAKLT CRISNVPYDHELKDMNVVWIRQRDNTELETRIGQAMEQENNELISVDVTATVCPEEWDSG ETFRCKVSIPSLLPTAEVRTLKKLKGGTPHAPAVYVLPPPSEQLALKETATLTCLMKDFY PNDFFVKWLRNDEPVADSEYFTSKPIQESKTPERYFTYSMLNVKEQDWNYGAIYTCVVGH EALPLLTTQKTLDKNT

#### >IgM4_Euleptes_europaea

TQNGPSLFPLILQSENLPSHGPITVGCLAKDVPPALVRFTWEDRSNVSVNATYIKQYPTV GNSAGTFTASSQLAISAEDWENFEPFYCKAESVDGVARVVRQASQTTEPRMIIRVPRCQD FENPNIKATVVCLADHLPTGSTTLQWLKDGNILDSGFTTTGPVANGCNGYSIISELPVSK QDWNSGKEFSCSVRNNMFSDTKNISKPTVCNSEVRVETIPPPFIDIYLSKSVNLTCRISN APDEEDLKALTVTWTRVSDKKNLETIIGQIKKEENREMFFVDATATICKEEWERGDTFEC KVTGSHLPTPEIKTLKKLHGFYPNDFFVKWLRNDEPVADSEYFTSKPIQESGTPERYFTY STLNVREQDWSYGAIYTCVVGHEALPFQTTQKTLDKNMGKPTLVNVSLVLSDTANTCY

#### >IgM1_Gekko_hokouensis

SQSAPSLFPLIPSDYTNAGDVTVGCLAKDFLPDSLTFSWQKPNNDSLEAEKIRRFPSIAN SNGVYTTSSEATIPARQWEDFEPYFCKAQLASETRVARVVRQKDFRGAYLNATLLCKADN LHTEKTTIKWLEDGNVLSSGFTTSAPIKQAGGGYSIISELIVTKRDWFANKIFSCQVQNE KYNEIRNVSKPSVCEIRVETIPPSFEDIYLTKSVKLTCRVSNIPFDEDLAALNVTWIRKH ANKPEEELETTIGQAKEQEDRELVFVDATANVCIEDWESGDEFQCKVALPLLASTETRSL RKENGRLFCIIPDPRNGGSRSPPSVYVSPPSSEELALKETATVTCLIKDFFPSGFFVRW LQNNEVVGDSEYYTSNPIQESKSPKRFFAYSTLRTTEQAWNEGTTFTCVVGHEALPYQTT QKTIDKNT

#### >IgM2_Gekko_hokouensis

SSRAPSLFPLIVPSESIPSNGLVTIGCLAKDVMPSVVSFTWNDQNNVSIDAHYVKQFPSI FYPSGTFTSSSQVAISANDWRNFQHFYCKATNNAGTGVVQVARQTPCLKPEMIIRAPRLE DFENSDPNATIVCMAANLHTARATVQWLKNGRTLNSGFANTGPVAVGPSGYSIISELTVD KRDWTSDKTFSCEVRNENFTSIRNISKSLVCDAGSCGEVQVHVETIPPSFADIYLNKSVK LTCRISNMPYDQELKELSVTWTRERDNTQLETVMGQITAQEDKALAFVDATATVCPEEWD SGDTFKCKVTIPSLLPTAETRTLRKLKDSPYHAPAVYVLPPPPEQLALQEVATLTCLMKG FYPNNFFVKWLRNDEPIGPSEYFISKPIQESKTPERYFTYSTLNVKEQDWSYGAIYTCVV GHEALPFQTTQKTIDKKT

#### >IgM3_Gekko_hokouensis

SPAAPSLFPLILPSESIPSNGLTTIGCLAKDVVPPVISFTWDNQNVSIDANNIKQFPSVF NPAGTFTSSSQVTISANDWWNFQPFHCTANNTNGTGVAQVIRPTTCLKPEMIIRAPRLED FDNSDPNATIVCMAVNLHTARATVKWLKKGHVLNSGFTTTGPAVMGRNGYSIVSELTVDR RDWVSEKTFSCEVHSENFTSIKNISKPLVCDLGPCSDVQVRVETIPPSFTDIYMTKSAKL TCRISNVPYDQELKELSVTWTRERDNTQLETVIGESIAQEDNEFIFVDATATVCPEEWDS GDIFKCKVTIPSLLPTAETRTLQKLQGGTPHAPAVYVLPPPSEQLTLQETATLTCMVKGF YPNEFFVKWLRNDEPLGDSEYFTSQPIQESKTPERYFTYSTLNVKEQDWSYGAIYTCVVG HEALPLQTTQKTIDKNT

#### >IgM4_Gekko_hokouensis

TQSGPSLFPLVIPSASIPSNGLMTIGCLAKDVLPALVSFSWSDYNNVSINAEYVTQYPTI GNSSGPYSSVSQVTVSAHDWRNFQPFFCKAEHVDGEAQVVRHSKAYLVSFDRQVTSPAS CTEPEMIIRAPRLEDFENSGPNATIVCMAANLHTKSITVQWLKDGHVLDGGVANTGPVAV GPSGYSIISELAVDRRDWISDKRFSCEVHNENFTSIKNISKSLVCDTGPCSDVQVHVETI PPSFTDIFMTKSAKLTCRISNMPYDQELKELSVTWTRERDNTQLETVIGESIAQEDNEFI FVDATATVCPEEWDSGDTFKCKVTIPSLLPTAETRTLQKPQGGTPRAPAVYVLPPPSEQL TLQETATLTCMVKGFYPNEFFVKWLRNDEPVGDSEYFISQPIQESKTPERYFTYSTLNVK EQDWSYGAIYTCVVGHEALPLQTTQKTIDKNT

#### >IgM5_Gekko_hokouensis

TQSGPSLFPLLLPCENTPSNGLVSIGCLAKDVPPALVRFTWEDNNNASINASYIKQYPTI KNSAGTFTASSQVTIPAGDWENSKPLYCKAESVVGVARVIPPASQLPAKMVIHAPRQIDV DNPDVNATISCMAAHLHTASTTIQWFKDGNILDSGFTTTGPLSVGSSGYSIFSELTVSKK DWSTGKIFSCRVHNEMFSNIKNISNSGCDPMEVHVETIPPSFIDIFLTKSAKLTCRISNL PYEEDQQELNVTWTRTSDKKELETIIGEAKKQENSDTFFVDATATICKEEWESEDTFECK VTHPLLPTAKTTTLTKLHGGTPSPPAVYVHPPTPEELALRETATVSCLLKSFYPKDAFVQ WLRNNELIEVGYFTSSPVQESKSPERYFTYSMLNINEQDWSNGDTYTCVVGHEALPLQTT QKTVDKNT

#### >IgM1_Gekko_kuhli

SQSPPSLFPLISSDYTNAGDITVGCLAKDFLPDSLTFSWQKPNNESLEAEKIRRYPSLAN SNGVYTTSSEATIPTNQWEASEPFFCKAEHSSGTRVARVVRPRNVLSSGFTTSAPTKQAH GGYSTMSELIVTKKDWFADKVFSCQVQNEKYDEIRNVSKLSVCEVRVEAIPPSFEDIYMT KSVKLTCRTSNIPFEQDLAELNVTWIRQHSNGGLEVLPTEIGQSKEQEDRDLVSVDATAN VCIEDWESGDNFQCIVALPLLASTETRSLKKENGGSRSPPSVYVQPPPLEELALKETATI TCLIKNFFPGDFFVRWMRNNEPVGVSESYTSNPMQESKSPKRYFSYSLLKVTEQEWSAGD TFTCMVGHEALPYQTTQKTIDKNT

#### >IgM2_Gekko_kuhli

PSKAPSLFPLIVPSESIPSNGLVTVGCLAKDVMPSVASFSWSDQNNVSIDAHYVKQFPSV FSPAGTFTSCSQVAISANDWRDFQPFYCKATNNAGTGVVRVTRQTSCLKPEMIIRAPRLE DFENSDPNATIVCMAANLHTARATVQWLKNGHTLDSGFTNTGPVAVGHSGYSIISELTVD KRDWTSDKTFSCEVRNENFTSMRNISKSLVCDTDNALAFVDATATVCPEEWDSGDTFKCK VTIPSLLPTAETRTLRKLKGRYFFDFTKKRINQPPQDSPYHAPTVYVFAPPPEQLALQET ATITCLLKGFYPNDFFVKWVRNDEPIGPSEYFISKPVQESKSPKLYFTYSTLNVKEQDWS YGATFTCVVGHEALPFQTTQKTIDKKT

#### >IgM3_Gekko_kuhli

PPAAPSLFPLILPSESIPSNGLTTIGCLAKDVVPPVISFTWDNQNVSINADNIKQFPSVF NPNGTFTSSSQVTISANDWWNFQAFHCVANNTKGTGVAQVIRPTSCLKPEMIIRAPRLED FENSDPNATIVCMAVNLQTARATVQWLKDGNVLKSGFATTGPAVMGRNGYSIVSELTVDR RDWVSEKMFSCEVHSENFTSIKNISKPLVCDLGSCSDVQTHVETIPPSFTDIYMTKSAKL TCRISNVPYDQDLKGLQVTWTREHDNTQLETIIGQATAQEDNTLVFIDATATVCPEEWDT GETFKCKVTFPSVLPTAETRTLQKLKGSTHYAPAVYVLPPPSEQLTLQETATITCLVKGF YPNDFFVKWLRNDEPVEDSQYLTSQPIRESKGPERYFTYSMLNVEEQDWSYGATYTCVVG HEALPLQTTQKTIDKKT

#### >IgM4_Gekko_kuhli

TQSGPSLFPLVMPSASIPSDGLMTIGCLAKDVLPALVSFSWNDYNNISINAQYVTQYPTI GNSSGPYTSVSQVTVSAHDWRNFQPFVCKAEHVDGEARVIRHSKTYLVSFDRQITSPTS CTEPEMIIRAPRLEDFENSGSNATIVCMAANLHTKSITVQWLKNGNTIDSDIANTGPVAV GASGYSIISELRVDRRDWFSDKTFSCEVHNENFTSIKNISKSLVCDTGSCSDVQVHVETI PPSFTDIFMTKSAKLTCRISNMPYDQELKELSVTWTRERDDTQLETIIGESIAQEDNEFI FVDATATVCPEEWYSGDTFKCKVTLPSLLPTAETRTLQKLQGGSPHVPAVYVLPPPSEQL TLQETATITCLVKGFYPNDFFVKWLRNDEPVGDSQYLTSQPIRESKSPERYFIYSMLDVK EQDWSNGDIYTCVVGHEALPLQTTQKTVDKNT

#### >IgM5_Gekko_kuhli

TQNGPSLFPLILPSENTPSNGLITFGCLAKDVPPSLVRFTWEDRNNVSINASYIKQYPTV KNSAGTYTASSQVAMSADDQEKFGPFYCKAESVVGRAWVDPQDFETPDVNATIFCMAAHL HTASTTIQWFKDGNILDSGFTTTRSVTVGCSGYSIFSELTVSKKDWTSGKEFSCRVHNEM FSNIMNISYTDCSGPVEVLVETIPPSFIDIYLTKSAKLTCRISNLPFKEDQKELNVTWTR ASDKKKLETIIGQPEKQENSGTFFVDATATICKEEWESEDTFECKVTHPLLPTAKTAVLT KLHGGTPSPPAVYVLPPPPEELALRETATLTCLLKGFYPKDTFVKWLRNNEPIEVGYFTS NPIQESKSPKQYSTYSMLTITEQDWSDGILYTCVVGHEALPFQTTQKTVDKKT

#### >IgM1_Heteronotia_binoei

SQTAPSLFPLIPSSDYTGDIVTIACLAKDFLPDSLTLSWQKPNNDSLEAEKIRRFPSIAN SNGVYTASSEATIPANQWEAFEPFFCKAELASQTRVTRVVRQKDFRSAYLNATLLCKADN LHTEKTTIKWLKNGQVLPSGFTTYAPVRGGRGSYSINSELIVTRKDWYAEEVFSCQVQND KFNLIKNVSKSSVCEGSTVPVRVETIPPAFADMFLTKTAYVTCRISNLPYGQDLAKLNVT WTREHSNGRVEILPTETGQPKEQENRELVYVDTTANICVEDWESGDNFQCTVRLPLLAST EIRNLKKQKGDSHNPPSVYVLPPPSEQLALQETATVTCLIRNYYPNDFFVQWMRNDEAVP ASDYYVGKPIEESKSPQRFFTYSTMNVNQQDWNYGARYTCVVGHEALPLNTIQKTIDKNT

#### >IgM2_Heteronotia_binoei

PQSAPSLFPLIVPSESIPSNGLVTVGCLAKDVMPDVVSFTWNDQNNVSIDANYVKQLPSV FNPSGTFTSSSQVAISANDWRNFEPFYCKATNKAGTGVVRVTRQTSCLKPEMIIRAPRLE DFENSDPNATIVCMAVNLQTDRATVQWLKDGHALNSGFATTGPAVMGRDGYRIISELTVD KRDWVSDKTFSCEVHNKNFTSIKNISKSLVCDTGPCTELKVQVKTIPPSFTDIYLTKSAK ITCRISNIPYDQDLKELEVIWTRERDNTQLETITGLATAQEDNAFIFVDATATVCPEEWD SGDTFKCKVTIPSLLPTAETRTLQKLKGSTYQAPAVYVLPPPSEQLDLQETATITCLIKG FYPNDFFVKWLRNDEPMGDSEYFISKPIQESKTPERYFTSSMLNVNEQDWSYGARYTCVV GHEALPLQTTQKTVDKNT

#### >IgM3_Heteronotia_binoei

SPAAPSLFPLILPSESIPSNGLMTIGCLAMDVVPPVISFTWDNQNVSINANNIKEFPSVF NPNGTFTSSSQVAISANDWWNFQPFYCKANNTIGTGVAQVIRPTSCFKPEMIIRAPRLED FVNSDPNATIVCMAVNLHTARATVQWLKDGHVLNSGFATTGPAVMGRNGYSIVSELIVDR RDWVSEKTFTCKVHSENFTSIKNISKPLVCDLGPCSDAQVHVKTIPPSFTDIYLTKSAKL TCRISNIPYDQDLQELKVIWTREHDNTQLETIIGQATAQEDNSLIFVDATATVCPEEWDS GETFKCKVTFPSVLPTAETRTLQKIKGRGSTYQAPAVYVLPPPSEQLALQETATVTCLVK SFYPNDFFVKWLRNDEPVGESEYFTIEPIQESKTPERFFTYSTLNVKEQDWNYGARYTCV VGHEALPLQTTQKTIDKNT

#### >IgM4_Heteronotia_binoei

TQSGPSLFPLIIPSASIPSNGLMTIGCLAKDVLPALVSFTWEDYNNININEKYVMQYPTI GNSSGTYSSVSQVTVSANDWRNFQPFYCKAEHVNGTARVIRHISCTEPEMIIRAPRLEDF ENSGRNATIVCMAANLYTTNTRVQWLKNGHALDSGFANTGPVAVGPRGYSVVSELIVDRR DWISDKTFSCEVRNENFTSIKNISKSLVCDTGPCDVQVHVETIPPSFTEIYLTKSAKLTC RISNIPYDQDLQELKVTWTREHDNTELETITGQATVQEDNSFIFVDATATVCPDEWDSGE TFKCKVIIPSLLPTAETRTLQKHKGSFYPNDFFVKWLRNDEPVGESEYFIIKPIQESKTP ERFFTYSTLNVKEQDWSYGARYTCVVGHEALPLHTTQKTIDKNTGKPTLVNVSLVLSEAT NTCY

#### >IgM1_Lepidodactylus_listeri

SQSAPSLFPLIPSDYTSVGDISVGCLAKDFLPDSLTISWQKPNNESLEAEKIRRFPSIAN SGGAYTTSSEATIPANQWEAFEPYYCKAQLASETRVARIVRQTPITCVDPTIIVRVPPLD DLRGPYLNATLLCKASNLHTEKTTLKWLKNGTVLSSGFTTSAPIKQAHGGYSIISELIVT RREWFSNEKFSCQVLNEKFDEIRNVSKSIACYVRVETIPPTFADIYLTKSAKLTCRISDI PFDQDLAQLNVTWIRQHANKPLEVLETEIGQTKEQDKELVFVDATTNVCVEDWNSGDTFQ CKVTFPALLASTQTRSLKKENGRLFYVIPDPQNGFYPNDFFVKWLQNGEPVGVAEYYTS NPIQESKPHKQYFTYSTLVISEQDWSSGTPYTCVVGHEALPFQTTQKTIDKNTGKPTLVN VSLVLSDVTSSCY

#### >IgM2_Lepidodactylus_listeri

SQSAPSLFPLIVPSESIPSNGLITVGCLAKDVFPSVNSFAWNDQNNVSINANYVKQFPSI FSPSGAFTSSSQVAISANDWWNFEPFYCKATNKAGTGVVQVARQTSCLKPDTIIRAPRLG DFENSDPNATIVCMAVNLHTARTTVQWLKNGHVLNSGFATTGPAVMGRDGYRIISELTVD KRDWESDKTFSCEVHNENFTSIKNISKSLVCDTGPSFDTQVHVETIPPSFVDIYMTKSVK LTCRISNMPYDQELKELNVTWIRGHDNTQLETIIGKATVQEDNKFVFADATATVCPEEWD SGETFKCKVAIPSLLPTAETRTLKKLTGSAHHAPAVYVLPPPLEQLALKETATLTCLVKK FYPNDLFVKWLRNDEPVADSEYFTSKPIQESKTPERYFTYSTLNVNEQDWSDGTTYTCVV GHEALPLQTTQKTIDKNT

#### >IgM3_Lepidodactylus_listeri

PPAAPSLFPLILPPESIPSNGLVTIGCLAKDVAPPVISFTWDNQNVSINADNVKQFPSVF NPAGTFTSSSQVAISANDWWNFQPFYCMANNTNGTGVAQVIRPTSCFKPEMIIRAPRVDD FENSDPNATIICMAANLHTARATVQWLKNGQVLNSGFATTGPAVMGRNGYSIISELTVDR RDWVSEKTFSCEVHSENFTSIKNISKLLVCGLGPCSDAQVHVETIPPTFTDIYMTKSAKL TCRISNMPYDQELKELSVTWTRERDNTQLETIIGEATPQEDNQFIFVDATAAVCPEEWNT GETFKCKVTFPSLLPTAETRTLKKLKGSTYYAPAVYVLPPSSEQLVLQETATITCLVKGF YPNDLFVKWLRNDEPVGDSEYFISQPIKESKTPERYFTYSMLNVKEQDWSSGVAYSCVVG HEELPLQTTQKTIDKNT

#### >IgM4_Lepidodactylus_listeri

TQSGPSLFPLIIPSESIPSNGLMTIGCLAKDVVPALVSFSWNDYNNISINAKYAMQYPTI GNSGTYSSVSQVTVSAHDWRNFQPFYCKAEHVDGEAQVVRQTSCTEPEMIIRAPRLEDFD SFGPNATIVCMAANLHTKSVTVQWLKNGHALDSGVANTGPVAVGPRGYSIISELAVDRRD WISDKTFSCEVHNENFTSIKNISKSLVCDTGSCTDVQVHVETIPPSFTDIYMTKSAKLTC RISNMPYDQELKELSVTWTRERDNTQLETIIGESTPQEDNEFVFADATATVCPEEWDSGE TFKCKVTIPSLLPTAETRTLQKPKGGTPRAPAVYVLPPPSEQLALQETATVTCLLKDFYPNEFFVKWLRNDEPVADSEYFISQPIQESKTPERYFTYGTLNVKEEDWSNGARYTCVVGHE ALPLQTTQKTVDKNT

#### >IgM5_Lepidodactylus_listeri

TQSGPSLFPLILPSENTPSSGLITIGCLAKDVPPALLRFTWEDHNNISINASYVKQYPTI KNSAGTFTASSQVSISADDWENLEPLYCKAESVVGAARVIRQEPKMVIHAPRRTDFENPD TNATISCKADHLHTASTTIQWLKDGKILDSGFTTTEPVAVGCSGYSIFSELTVTKKDWST EKLFSCRVHNEMFSNIKNISSSMTCDPGSGGNVEIRVETIPPSFFDIYMTKSAKLTCRIS NIPFEEDLKQLNVTWTRESDKKALETIIGQVKKQENSDLFFVDATATICKEEWESKDTFE CKVTHPHLPTAKTTILRKLHGETPHAPAVYVLPPPSEQLALRETATITCLLKGFYPKDIF VKWLRNNEPIQVGYFTSNPVQESKSPERYFIYSILNINEQDWSDGTPYTCVVGHEALPFQ TTQKTVDKNM

#### >IgM1_Lepidodactylus_lugubris

SQSAPSLFPLISSDYASAEDITVGCLAKDFLPDSLIFSWQKPNNESLEAEKIRRFPSISN SDGAYTTCSEATIPANQWDAYKPFFCKAQLASETRVARVVRQTPSICEEPTIIVRVPPLD DFQGAYLNATLLCKADNLHTEKTTLKWLKDGIVLSSGFTTSAPIKQSRGGYSIISELIVT KREWFANKVFSCQVQNEKFDEIRNASKPLVCEVRVETIPPAFADIYVTKSAKLTCRTSNI PFTEDLADLNVTWIRQPTNKPVEILETNIGQAKEQEDRELVFVDATTTVCVEDWNSGDTF QCKVTFHQLASTETRTLKKENGRLFVIPDPQNGGSRSPPSVYVLPPSSEQLALKETAT LTCLVKDFYPNSFFVQWLQNNELVGVSEYHTSDPIQESKSPERYFTYSTLSVSEQDWSSG ATYTCVVGHEALPLQTTQKTIDKNT

#### >IgM2_Lepidodactylus_lugubris

SQSAPSLFPLIVPSESIPSNGLVTVGCLAKDVMPSVDSFAWNDQNNVSINANYIKQFPSV FGPSGTFTSSSQVSISANDWRNFEPFYCKATNKAGTGVVQVARQTSCLKPATIIRAPRLE DFENSDPNATIVCMAVNLQTARTTVQWLKNGHVLDSGFATTGPAVMGRDGYRIISELTVD KRDWESDKKFSCEVHNENFTSIKNISKSLVCDTGPSFEAQVHVETIPPSFIDIYMTKSVK LTCRISNMPYDQELKELNVTWTRGRDNTQLETIIGEATVQEDDKFIFADATATVCPEEWD SSAYHPPAVYVLPPPSEQLALKETATITCLLKGFYPNDCFVKWLRNDEPVGDSEYFTSQP IQESKTPERYFTYSTLNVNEQDWSYGATYTCMVGHEALPLQTTQKTIDKNT

#### >IgM3_Lepidodactylus_lugubris

PPAAPSLFPLILSPESIPSNGLVTIGCLAKDVVPPVISFTWDNQNVSINADNVKQFPSVF NPAGTFTSSSQVAISANDWWNFQPFYCMANNTNGTGVAQVLRPTSCFKPEMIIRAPRLED FKNFNPNATIVCMAANLHTARATVQWLKKGHVLNSSFSTTGPAVMGHNGYSIISELTVDR RDWVSDKIFTCEVHSENFTSIKNISKQLVCDLGPSFEAKAHVETIPPTFTDIYMTKSAKL TCRISNIPYDQELKQLQVTWTRERDNTQLETITGEAMAQEDDQFVFVDITATVCPEEWNT GETFKCKVTFPSILPTAETRTLQKLKGSTYYAPAVYVLPPPSEQLALQETATITCLLKGF YPNDFFVKWLRNDEPVGDSKYFTSKPIQESKTPERYFTYSTLNVKEQDWSYGATYTCVVG HEELPLQTTQKTIDKNT

#### >IgM4_Lepidodactylus_lugubris

TQSGPSLFPLIIPSESIPSNGLMTIGCLAKDVVPALVSFSWNDYNNISINAKYVMQYPTI GNSSGMYSSVSQVTVSAHDWRNFQPFYCKAEHVDGEAQIVRRSKTFFLSFDPLIASPTS CTEPEMIIRAPRLEDFESSGPNATIVCMAANLHTKSVTVQWLKNGLALDSGVANTGPVAV GPRGYSIISELIVDRRDWISDKTFSCEVHNENFTSIKNISKSLVCDMGPCTDVQVQVETI PPTFTDIYMTKSVKLTCRISNMPYDQELKELQVTWTRGRDNTQLETIIGEAMPQEDNEFV FADATATVCPEEWDSGETFKCKVTIPSLLPTAETRTLQKPKGGTPRAPAVYVLPPPSEQL ALQETATLTCLLKGFYPNEFLVKWLRNDEPVGDSEYFISKPIQESKTPERYFTYSTLNIK EQDWSYGVTYTCMVGHEALPLQTTQKTVDKNT

#### >IgM5_Lepidodactylus_lugubris

TQSGPSLFPLILRSENTPSSGLITIGCLAKDVPPALLRFTWEDHNNVSINASYVKQYPTI KNSAGTFTASSQVAISADDWENFEPFYCKAESVVGAARVIRQDFEKPGINATISCKADHL HTANTTIQWLKDGNILDSGFTTAGPIAVGCSGYSIFSELTVTKKDWRTETLFSCRVHNEM FSDIKNISSSMTCDSGCGDDVEVHVETIPPSFIDIYQTKSAKLTCRISNMPYKEDLTELN VTWTRESDKKALETIIGQVKKQENSDLFFVDATATICKEEWESKDTFECKVTHPLLPAAK TTILRKLHGGTPQAPAVYVLPPPSEQLALRETVTLTCLVKGFYPNDFFVKWLRNDEPIGV GYFTSNPVQESKSPERYFIYSMLNINEQDWSDGTSYTCMVGHEALPFQTTQKTIDKNT

#### >IgM1_Mediodactylus_kotschyi

SQTAPSLFPLIPSSDNTISGDAITVGCLAKDFLPDSLTLSWQKPNNESLEAEKVRRFPSF ANSNGIYSACSEATIPADQWKAYEPFFCKAELASQTKVARVVRQTPIVCLPPEIMIRVPP LEDFRGPYLNATILCKASNLHTEKTTIKWLKEGKVLDSGFTTSAPIKQLRGGYSLVSELI VTRKDWYSDNIFSCQMENDKFNEIRNVSKPGVCEVRVETIPPSFADMYLTNSAKLTCRIS NIPYSEDLAELNVTWIREHSNKRIEVLETVIGQAEEQENGDLLFVDATTNVCVEDWETGD NFQCKVRLPLLASTEIKSLRKQNETATLTCLLNNFYPNDMFVKWVRNDEPIGESEYFTSK PIQESKTPERYFAYSMLKVKEQDWSYGATYTCVVGHEALPYKTTQKTIDKNT

#### >IgM2_Mediodactylus_kotschyi

SPTAPSLSPLILPSESIPNNGLMTIGCLAKDVVPPVISFSWDYQNVSINANTIKQFPSVF DPAGTFTTSSQVTISANDWWNFQPFHCMANNTIGTGMAQVIRPTSCLKPEMVIHAPRLED FENSDRNATIVCMAVNLHTTSTTVQWMKNGHALDSGFTTTGPAVMGRSGYRIISELTVDR RDWVSDKIFSCEVHNENFTSIKNISKPLVCDLGPCSDAQVHVETIPPTFTDIYMTKSAKL TCRISNIPYDQDLKELKVTWTKEHDSTELETTTGQATAQKDDQWIFVDATASVCPEEWNT GETFKCKVIIPSLLPTAETKTLQKLKGGTPSPPSVYVLPPPSEQLALQETATITCLLKGF YPNDFFVKWLRNDEPIADSEYFISEPIQESKTPKRYFTYSTLNIKEQDWSYGTTYTCVVG HEALPLQTTQKTIDKNT

#### >IgM3_Mediodactylus_kotschyi

TQSGPSLFPLILPSASIPSNGLMTIGCLAKDVLPAFSSFSWNDYNNISINAKYVTQYPTV GNSSGAYSSISQVTVSAHDWRNFQPFYCKAEFVDGAARVVRHSKTHLISFDQLITSLTS CTEPEMIIRAPRLEDFETSGPNATIVCMAVHLHTTSVTVQWLKNGRALNSGVANTGPVAV GPSGYSIISELTVDRRDWVSDKTFSCEVHNENFTSIKNISKSLVCDLGPCTDVQVHVETI PPTFTDIYMTKSAKLTCRISNIPYDQDLKELKVTWTKEHDNTELETTIGQATAQKDDQWI FVDATASVCPEEWNTGETFKCKVIIPSLLPTAETKNLQKLKVGSPRPPSVYVLPPPSEQL ALQETATITCLLKGFYPNDFFVKWLRNDEPIADSEYFISEPIQESKTPKRYFTYSTLNIK EQDWSYGTTYTCVVGHEALPLQTTQKTIDKNT

#### >IgM4_Mediodactylus_kotschyi

NQSGPSLFPLILPSENTPSNGLVTIGCLAKDVPSALVRFTWQDHNNVSINASYIKQYPTI STSPGTFTASSQVAVSADDWQKFEPFYCKAESIVGSARVIRQSKISPHTKPAMTIHAPR RADFEDPTVNATISCTAAHLHTTSTTIQWLKDGNILDSGFITTGPATAGCGYSIISELTV SKKDWSADKEFSCKVHNEMFSDIKNISYSSICDSVHVETIPPSFIDIYLNKSAKLTCRIS NVPYKEDLKELKVTWTRASNKKELETIIGQAKNQENSDKFFVDATATVCKDEWESEDSFE CKVTLPLLPTAETKILRKLRGGTPSPPSVYVLPPPSEELALRETATITCLLKRFYPNDFF VKWLRNDEPIADSEYFTSKPVQESMSPEWYFAYSMLSVKEQDWSYGATYTCMVGHEALPL QTTQKTIDKNT

#### >IgM1_Paroedura_picta

SSSAPSLFPLIPSSDNTISGDSVIIGCLAKDFLPDSVSFSWQKPNNSSLEAGKIRSFPSI ANSNGIYTTSSEATVPADQWEATEPFYCKAEHNSQTRVARVVRPYCPPPTDPKITIHVPP LVEFQGPYLNATLLCKAENMHTERTTIKWLKDGIVLDSGFATTAPIRQHRGGYSITSELI VTKRDWYADKKFSCQVQNEKFNEMRNVSMSSTCESGGESNLRVIVQTIPPSFDDIFHNKA KLTCRISNLPYGQDPAEVNATWIRERNNEVLHTELGQTKEQENKELVYVDATAEICREEW DSGDNYLCIVEIPSLLATTERKSLKKQNDFYPNDFFVKWMRNSKPVEGSEYWTSKPMQES KSPERYFTYSTLSIPEQDWSSGDTYTCVVGHEALPYQTTQKTIDRNTGKPTLVNVTLMLS DVTSTCS

#### >IgM2_Paroedura_picta

PPAAPSVFPLTLPPESIASSGLVAIGCLAKDVVPHIISFSWDNRNVSINAENVKQFPSVF HPAGTFTSSSQVAISANDWWNFQPFYCTANNSNGTGVAQVVRPTSCLKPEMIIRAPRLED FKSSDPNATIVCMAVNLHTARATVQWRKNGQALSSGFTTSGPAVMGRDGYSIVSELTVDR RDWISEKTFSCEVHSENFTSIKNISKPLVCDLGPCSDVRVDVKTIPPSFNDIFTTKSAKL TCRISNMPYDQDLKDLEVVWTRERDNTELKTTIGEATAQENDELIFVDATATVCPEEWDS EDTFKCKVTFPSVLPTTETRTLKKLLSFYPNDFFVKWMRNNEPVGDAEYFISEPVQESKS PERYFTYSAFNVPEQDWSYGALYTCVVGHEALPLQTTQKTIDKNTGKPALVNVSLVLSEA TQTCY

#### >IgM3_Paroedura_picta

TQSGPSLFPLIIPSASIPSNGLMTIGCLAKDVPPALVSFSWDDYNNVGISAKYVTQYPTI GNSSGGFSAVSQVTVSAHDWRNFQPFYCKAEHVNGTGRVVRHSKILFIYLYFYLYTAAL TKTPSCTEPEMIIRAPRLGDFENPGTNATIVCMAANLHTASVTVHWLKNGHALDSDFANT GPVAVGPSGYSVISELTVDRRDWVSDKMFSCEVHNENFTSIRNISKNLVCDAGPCSDVRV DVKTIPPSFNDIFTTKSAKLTCRISNMPYDQDLKDLEVVWTRERDNTELKTTIGEATAQE NDELIFVDATATVCPEEWDSEDTFKCKVTFPSVLPTTETRTLKKLLSFYPNDFFVKWMRN NEPVGDAEYFISEPMQESKSPERYFTYSAFNVPEQDWSYGAIYTCVVGHEALPLQTTQKT IDKNTGKPTLVNVSLVLSEATQTCY

#### >IgM1_Sphaerodactylus_townsendi

TQTAPGLFPLIPSSDSYTNGDVTIGCLAKDFLPDSLTFSWQNPNNDSLEASKIRQFPSIM KSTGVFITSSEATIPASQWNTFEPFFCKAKHPSETKVTRVVRQTPIVCLPPKVIVRVPPL EDFKGPYLNATLFCKASGLHTERTTITWMKSENVLTSGFTTTAAVREGRGSFSITSELIV TKKDWYANKVFSCQVQNEKFNEMRNVSMLTVCEVETIPPSFGDMYLNNTVTLTCRISNLP FGLGLDELNVTWIRERGSTPLNTSTGQPEDQENSELVFVDATATVCREEWENGDNYLCKV RHPQLASTKTKSLKKQKGRLFYIIPQNWRTSLPPSGSRSPPSVYVLPPPAEQLALRET ATVTCLLKDFYPNDIFVRWMRNNEPVGASEYYISQPIRESKSPERYFAYSTLNVKEQDWS YGATYTCVVGHEALPLQTTQKTVDKST

#### >IgM2_Sphaerodactylus_townsendi

SPGPPSLFPLILRPESIPSSGLVTIGCLAKDVVPPVISFTWDHQNVSINPDSIKQFPSVF NPVGTFSSSSQVVISAHDWWNFQPFYCRANNTNGNGVVQVVRPTSCLQPEMIIRAPRLED FENSNPNATIVCMAVNLRTARATVQWQKNGHVLNSGFATTGPAVMGRNGYSIVSELIVDR RDWVSEKTFSCEVRSENFTSTKNISKPLVCDIGPGSEMKVHVGTIPPTFTDIYLTKSAKL TCRISNIPYDQVPKKLDVIWTRERDNKELETKIGQATAQDKNELVFVDATAAVCPEEWNS GETFKCRVTIPSLLPATETRILQKLKSSAYHPPAVYVLPPPSEQLALQETATITCLLKGF YPNDFFVKWMRNNEPVGPSEYFTSQPIQESKTPERYFTYSTLNVKEQDWSYGDTYTCVVG HEALPLQTTQKTVDKNT

#### >IgM3_Sphaerodactylus_townsendi

TQSGPSLFPLILPSEKIPRNGPVTVGCLAKDVPPALVRFTWEDHNNGSINSSSIKQYPTV GNSAGTFTASSQMTVSAEDWENFQPFYCKAENVNGSARIVRRAAQTEPKLIIRGPRQQDF KNPNKNANIVCVGSHMHDASTTVQWLKDGNILDSGFTTTGSAAAGCGGYSVVSELAVSKA DWSADKKFSCRVHNKMFSRTENISSSMICDSGDIVEVHVETIPPSFVDIYLTKSANLTCR ISNLPEEEDLKELHVTWTRVSDKKELDTVIGQVKRQENSEMFFVDATATICKEEWDKDAF ECKVTHHLLPAAEIKTLRKLHGGTPHAPAVYVLPPPSEQLALQETATITCLVKGFYPNDF FVKWMRNDEPVGPSEYFTSQPIQESKTPERYFAYSTLNVKEQDWSYGATYTCVVGHEAFP LQMSQKTLDKNT

#### >IgM1_Teratoscincus_roborowskii

SPSAPSLFPLIPSDITSSGDITVGCLAKDFLPDSLTLSWQKPNNDSLEAAKVRQFPSVLK SNGVYTTSSEAAVPAAQWTANEPFYCKAELSSQTTVARVVRANFRGPYLNATLFCVADHL HTERTTIKWLEGGNVLTSGFTTTAPIRQDGGYSITSKLIVSKRDWYADKTFSCQVQNDKF NEIRNVSRLSVCESLYPPSAAVHVETIPPSFADIYLTKSAKLTCRISNIPYSEDLEELNV TWIRERDNKPLETTIGGSHSPPSVYVLPPPSEQLALRETATLTCLLKDYYPKDSFVKWMR NDEPVGATEYFTSTPVQESKSPERYSSYSMLNVNEQEWNSGDTFTCVVGHEALPLQTTQK TVDKNT

#### >IgM2_Teratoscincus_roborowskii

NQTGPSLFPLISPYESIPSNGLMTIGCLAKDVPPALVSFSWDDYNNVSINAKYITQYLTI GNSAGTFSSVSQVAISANDWRNFQPFYCKAENVNGAAQVVRHSKTSLVSFDQPTSSLEP QPPCIRPEVIIRAPRLEDFENSGPNATLVCMAVNLHTTSTKVQWLKNGRAVDSDFTTTGP VAVGPSGYSIISELTVDRRDWVSDKTFSCEVHNENFTSIKNISKSLVCDLGSCSDVQVQV ETIPPSFTDIYMTKSAKLTCRISNVPYDQDLKELNVIWTRERDNTQLETRIGQYTAQEKN ELVFVDATATVCPEEWDSGETFKCKVTIPSLLPTAEERTLKKHQGGTPRAPAVYVLPPSS EQLALQETATITCLLKGFYPNDFFVKWLWDDEPVGASEYFTSKPIQESKNPERYFTYSTL NVKEQDWSNGATYTCVVGHEALPLQTTQKTVDKNT

*Figure S1. Three-dimensional models of IgM1, IgM4 and IgM5 from Eublepharis macularius*

**Figure 6:**
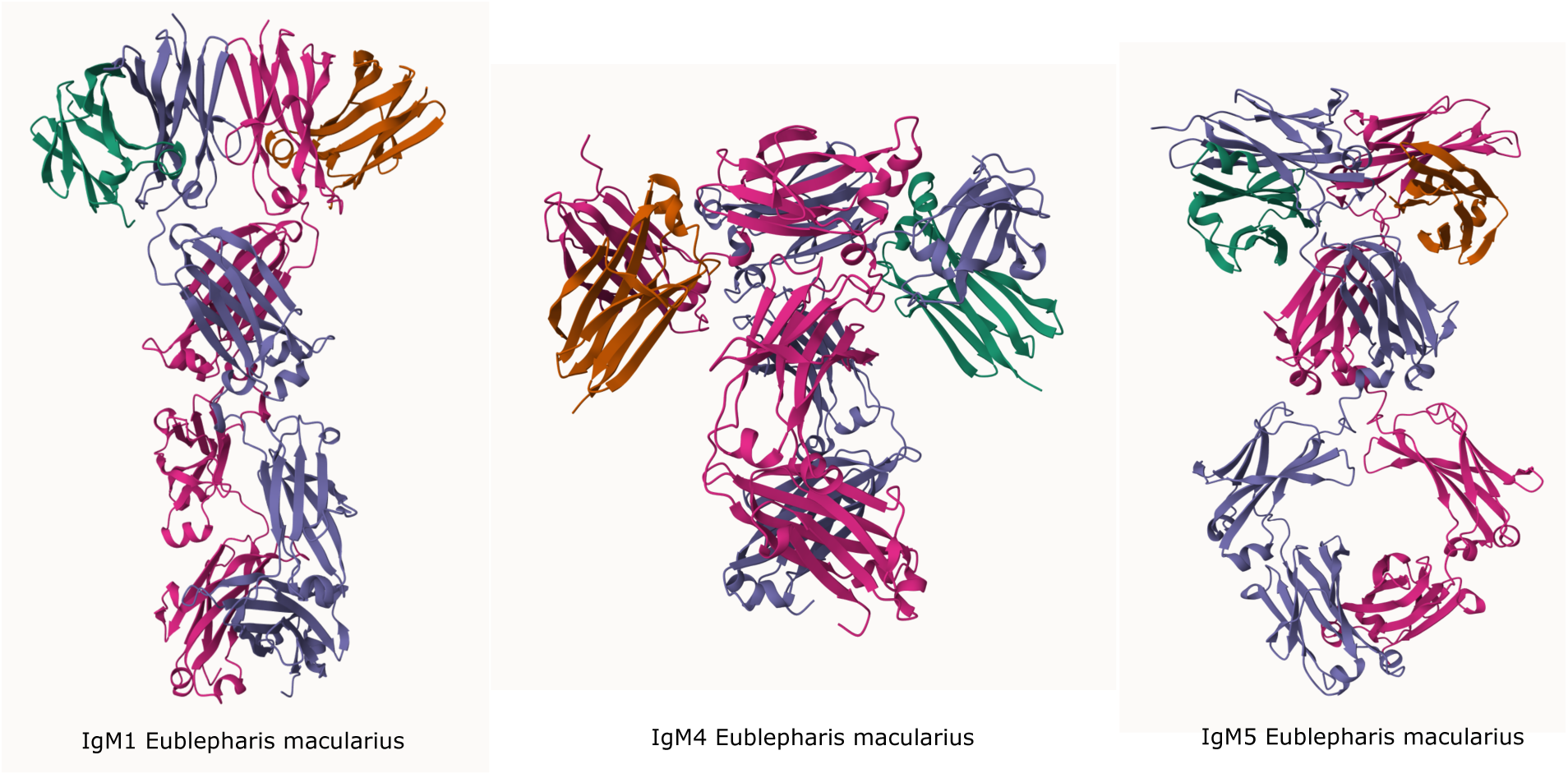
Three-dimensional models of IgM variants from *Eublepharis macularius*. The figure shows the 3D representations of IgM1, IgM4, and IgM5 analyzed in detail in this study.

**Table S1.** Complete list of diagnostic positions.

| Position | Domain | IgM1 aa | Freq. | Other aa | Freq. | Change type |
| --- | --- | --- | --- | --- | --- | --- |
| 12 | CH1 | P | 85% | L | 59% | hydrophobicity |
| 13 | CH1 | S | 100% | P | 79% | conservative |
| 15 | CH1 | D | 83% | E | 82% | conservative |
| 22 | CH1 | D | 62% | L | 85% | charge, hydrophobicity |
| 32 | CH1 | F | 100% | V | 95% | size |
| 35 | CH1 | D | 92% | A | 51% | charge, hydrophobicity |
| 37 | CH1 | L | 92% | V | 54% | conservative |
| 38 | CH1 | T | 77% | S | 67% | conservative |
| 43 | CH1 | K | 92% | D | 74% | charge |
| 49 | CH1 | L | 100% | I | 87% | conservative |
| 50 | CH1 | E | 100% | N | 79% | charge, polarity |
| 53 | CH1 | K | 85% | Y | 67% | charge, hydrophobicity |
| 55 | CH1 | R | 85% | K | 67% | conservative |
| 56 | CH1 | R | 69% | Q | 90% | charge |
| 72 | CH1 | E | 92% | Q | 87% | charge |
| 73 | CH1 | A | 100% | V | 90% | hydrophobicity |
| 76 | CH1 | P | 100% | S | 90% | polarity |
| 79 | CH1 | Q | 100% | D | 92% | charge |
| 82 | CH1 | A | 77% | N | 85% | hydrophobicity, polarity |
| 113 | CH2 | I | 75% | M | 77% | hydrophobicity |
| 117 | CH2 | V | 100% | A | 86% | hydrophobicity |
| 119 | CH2 | P | 100% | R | 89% | charge, hydrophobicity |
| 125 | CH2 | G | 92% | N | 67% | hydrophobicity, polarity |
| 128 | CH2 | L | 92% | P | 56% | hydrophobicity |
| 132 | CH2 | L | 75% | I | 87% | conservative |
| 133 | CH2 | L | 75% | V | 64% | conservative |
| 135 | CH2 | K | 92% | M | 74% | charge, hydrophobicity |
| 142 | CH2 | E | 83% | A | 54% | charge, hydrophobicity |
| 146 | CH2 | I | 75% | V | 67% | conservative |
| 147 | CH2 | K | 92% | Q | 90% | charge |
| 163 | CH2 | A | 92% | G | 87% | hydrophobicity |
| 170 | CH2 | G | 77% | S | 53% | polarity |
| 179 | CH2 | I | 92% | T | 67% | hydrophobicity, polarity |
| 181 | CH2 | T | 92% | D | 72% | charge, hydrophobicity |
| 187 | CH2 | A | 77% | S | 69% | hydrophobicity, polarity |
| 194 | CH2 | Q | 100% | E | 67% | charge |
| 196 | CH2 | Q | 77% | H | 77% | conservative |
| 199 | CH2 | K | 100% | N | 62% | charge, polarity |
| 201 | CH2 | N | 77% | T | 62% | hydrophobicity |
| 202 | CH2 | E | 85% | S | 62% | charge, hydrophobicity |
| 204 | CH2 | R | 92% | K | 74% | conservative |
| 206 | CH2 | V | 92% | I | 95% | conservative |
| 210 | CH2 | S | 69% | L | 76% | hydrophobicity, polarity |
| 213 | CH2 | E | 85% | D | 89% | conservative |
| 223 | CH3 | R | 67% | H | 71% | charge |
| 256 | CH3 | A | 69% | K | 68% | charge, hydrophobicity |
| 317 | CH3 | A | 92% | P | 95% | hydrophobicity |
| 318 | CH3 | S | 75% | T | 82% | conservative |
| 319 | CH3 | T | 92% | A | 82% | hydrophobicity, polarity |
| 323 | CH3 | S | 67% | T | 68% | conservative |
| 331 | CH4 | S | 80% | T | 74% | conservative |
| 334 | CH4 | P | 80% | A | 74% | hydrophobicity |
| 336 | CH4 | S | 82% | A | 74% | hydrophobicity, polarity |

