## Supplementary material for "Evolutionary Divergence and Structural Differentiation of Multiple Immunoglobulin M Genes in Gekkota (Squamata: Reptilia)": latex: references.bib

% Revised references (checked/corrected where needed)
% Notes:
% - Flajnik & Kasahara year corrected to 2010.
% - Das et al. authors corrected (Nei instead of Bhattacharya) and DOI fixed.
% - Chen et al. author list corrected (no Bhagat) and DOI added.
% - Shannon 1948 is in two parts; kept as a single entry with both page ranges and both DOIs in note.

@article{bengten2006,
  title   = {Channel catfish immunoglobulins: repertoire and expression},
  author  = {Bengt{\'e}n, Eva and Clem, L. William and Miller, Norman W. and Warr, Gregory W. and Wilson, Melanie},
  journal = {Developmental and Comparative Immunology},
  year    = {2006},
  volume  = {30},
  number  = {1-2},
  pages   = {77--92},
  doi     = {10.1016/j.dci.2005.06.016}
}

@article{boes2000,
  title   = {Role of natural and immune {IgM} antibodies in immune responses},
  author  = {Boes, Marianne},
  journal = {Molecular Immunology},
  year    = {2000},
  volume  = {37},
  number  = {18},
  pages   = {1141--1149},
  doi     = {10.1016/S0161-5890(01)00025-6}
}

@article{chen2009,
  title   = {Immunoglobulin {D} enhances immune surveillance by activating antimicrobial, proinflammatory and {B} cell-stimulating programs in basophils},
  author  = {Chen, Kang and Xu, Weifeng and Wilson, Melanie and He, Bing and Miller, Norman W. and Bengt{\'e}n, Eva and Edholm, Eva-Stina and Santini, Paul A. and Rath, Poonam and Chiu, April and Cattalini, Marco and Litzman, Jiri and Bussel, James B. and Huang, Bihui and Meini, Antonella and Riesbeck, Kristian and Cunningham-Rundles, Charlotte and Plebani, Alessandro and Cerutti, Andrea},
  journal = {Nature Immunology},
  year    = {2009},
  volume  = {10},
  number  = {8},
  pages   = {889--898},
  doi     = {10.1038/ni.1748}
}

@article{cock2009,
  title   = {Biopython: freely available {Python} tools for computational molecular biology and bioinformatics},
  author  = {Cock, Peter J. A. and Antao, Tiago and Chang, Jeffrey T. and Chapman, Brad A. and Cox, Cymon J. and Dalke, Andrew and Friedberg, Iddo and Hamelryck, Thomas and Kauff, Frank and Wilczynski, Bartek and de Hoon, Michiel J. L.},
  journal = {Bioinformatics},
  year    = {2009},
  volume  = {25},
  number  = {11},
  pages   = {1422--1423},
  doi     = {10.1093/bioinformatics/btp163}
}

@article{das2008,
  title   = {Evolutionary redefinition of immunoglobulin light chain isotypes in tetrapods using molecular markers},
  author  = {Das, Sabyasachi and Nikolaidis, Nikolas and Klein, Jan and Nei, Masatoshi},
  journal = {Proceedings of the National Academy of Sciences of the United States of America},
  year    = {2008},
  volume  = {105},
  number  = {43},
  pages   = {16647--16652},
  doi     = {10.1073/pnas.0808800105}
}

@article{flajnik2010,
  title   = {Origin and evolution of the adaptive immune system: genetic events and selective pressures},
  author  = {Flajnik, Martin F. and Kasahara, Masanori},
  journal = {Nature Reviews Genetics},
  year    = {2010},
  volume  = {11},
  number  = {1},
  pages   = {47--59},
  doi     = {10.1038/nrg2703}
}

@article{gamble2012,
  title   = {Repeated origin and loss of adhesive toepads in geckos},
  author  = {Gamble, Todd and Greenbaum, Eli and Jackman, Todd R. and Russell, Anthony P. and Bauer, Aaron M.},
  journal = {PLoS ONE},
  year    = {2012},
  volume  = {7},
  number  = {6},
  pages   = {e39429},
  doi     = {10.1371/journal.pone.0039429}
}

@article{gambondeza2009anolis,
  title   = {The immunoglobulin heavy chain locus in the reptile {Anolis carolinensis}},
  author  = {Gamb{\'o}n-Deza, Francisco and S{\'a}nchez-Espinel, Christian and Magad{\'a}n-Momp{\'o}, Susana},
  journal = {Molecular Immunology},
  year    = {2009},
  volume  = {46},
  number  = {8-9},
  pages   = {1679--1687},
  doi     = {10.1016/j.molimm.2009.02.019}
}

@article{gambondeza2018shinisaurus,
  title   = {Immunoglobulin and {T} cell receptor genes in Chinese crocodile lizard {Shinisaurus crocodilurus}},
  author  = {Gamb{\'o}n-Deza, Francisco and Olivieri, David N.},
  journal = {Molecular Immunology},
  year    = {2018},
  volume  = {102},
  pages   = {81--93},
  doi     = {10.1016/j.molimm.2018.06.263}
}

@article{olivieri2021insights,
  title     = {Insights into the evolution of {IG} genes in amphibians and reptiles},
  author    = {Olivieri, David N and Mirete-Bachiller, Serafin and Gambon-Deza, Francisco},
  journal   = {Developmental and Comparative Immunology},
  volume    = {114},
  pages     = {103868},
  year      = {2021},
  publisher = {Elsevier}
}

@article{henikoff1992,
  title   = {Amino acid substitution matrices from protein blocks},
  author  = {Henikoff, Steven and Henikoff, Jorja G.},
  journal = {Proceedings of the National Academy of Sciences of the United States of America},
  year    = {1992},
  volume  = {89},
  number  = {22},
  pages   = {10915--10919},
  doi     = {10.1073/pnas.89.22.10915}
}

@article{katoh2013,
  title   = {{MAFFT} multiple sequence alignment software version 7: improvements in performance and usability},
  author  = {Katoh, Kazutaka and Standley, Daron M.},
  journal = {Molecular Biology and Evolution},
  year    = {2013},
  volume  = {30},
  number  = {4},
  pages   = {772--780},
  doi     = {10.1093/molbev/mst010}
}

@article{kyte1982,
  title   = {A simple method for displaying the hydropathic character of a protein},
  author  = {Kyte, Jack and Doolittle, Russell F.},
  journal = {Journal of Molecular Biology},
  year    = {1982},
  volume  = {157},
  number  = {1},
  pages   = {105--132},
  doi     = {10.1016/0022-2836(82)90515-0}
}

@article{nei2005,
  title   = {Concerted and birth-and-death evolution of multigene families},
  author  = {Nei, Masatoshi and Rooney, Alejandro P.},
  journal = {Annual Review of Genetics},
  year    = {2005},
  volume  = {39},
  pages   = {121--152},
  doi     = {10.1146/annurev.genet.39.073003.112240}
}

@article{nimmerjahn2008,
  title   = {{Fc}$\gamma$ receptors as regulators of immune responses},
  author  = {Nimmerjahn, Falk and Ravetch, Jeffrey V.},
  journal = {Nature Reviews Immunology},
  year    = {2008},
  volume  = {8},
  number  = {1},
  pages   = {34--47},
  doi     = {10.1038/nri2206}
}

@book{ohno1970,
  title     = {Evolution by Gene Duplication},
  author    = {Ohno, Susumu},
  publisher = {Springer-Verlag},
  address   = {New York},
  year      = {1970}
}

@article{saitou1987,
  title   = {The neighbor-joining method: a new method for reconstructing phylogenetic trees},
  author  = {Saitou, Naruya and Nei, Masatoshi},
  journal = {Molecular Biology and Evolution},
  year    = {1987},
  volume  = {4},
  number  = {4},
  pages   = {406--425},
  doi     = {10.1093/oxfordjournals.molbev.a040454}
}

@article{shannon1948,
  title   = {A Mathematical Theory of Communication},
  author  = {Shannon, Claude E.},
  journal = {Bell System Technical Journal},
  year    = {1948},
  volume  = {27},
  pages   = {379--423, 623--656},
  note    = {Published in two parts: Part I DOI 10.1002/j.1538-7305.1948.tb01338.x; Part II DOI 10.1002/j.1538-7305.1948.tb00917.x}
}
