## Supplementary figures and images for "Evolutionary Divergence and Structural Differentiation of Multiple Immunoglobulin M Genes in Gekkota (Squamata: Reptilia)"

### 3D_Eublpharis.pdf

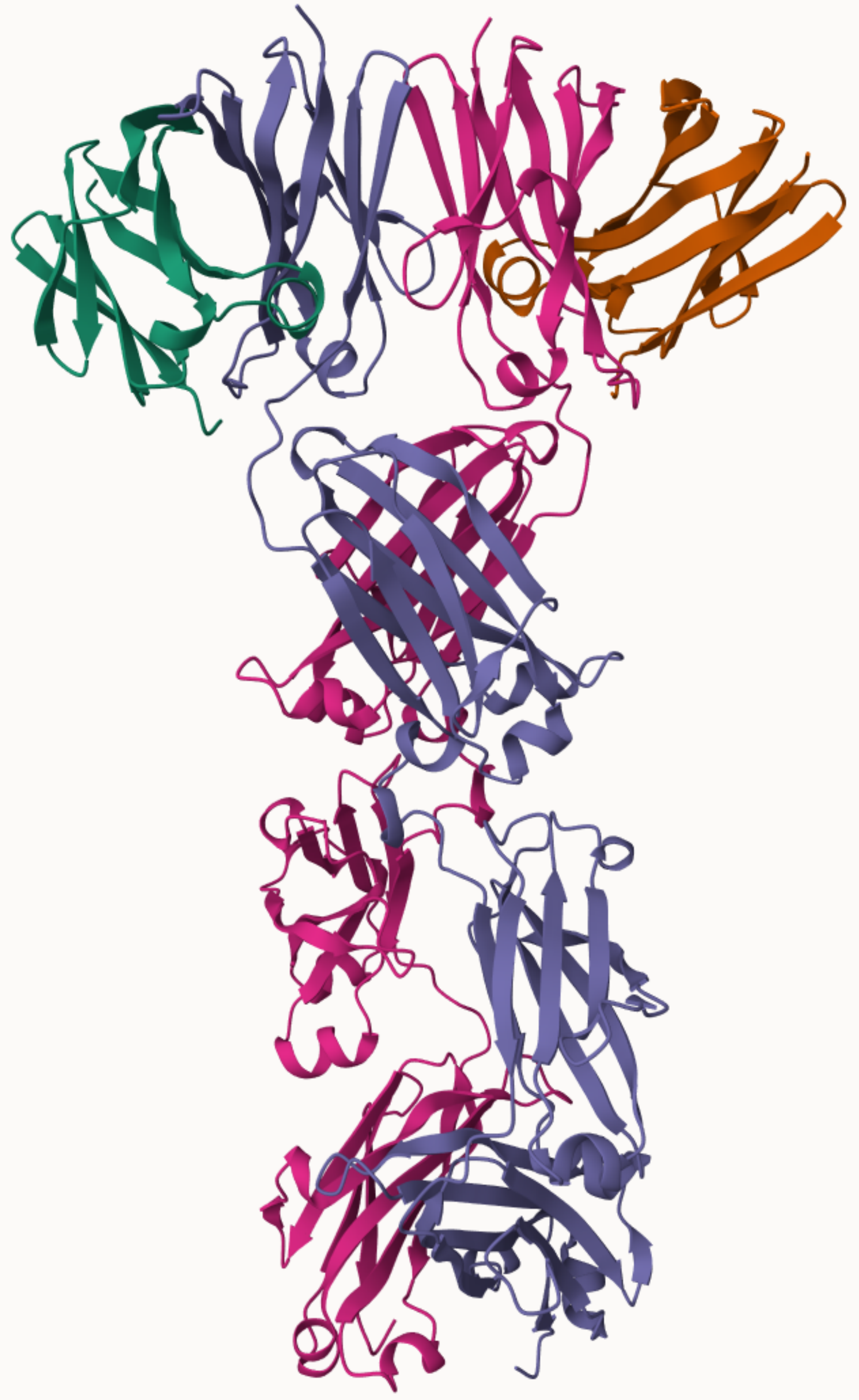

IgM1 Eublepharis macularius

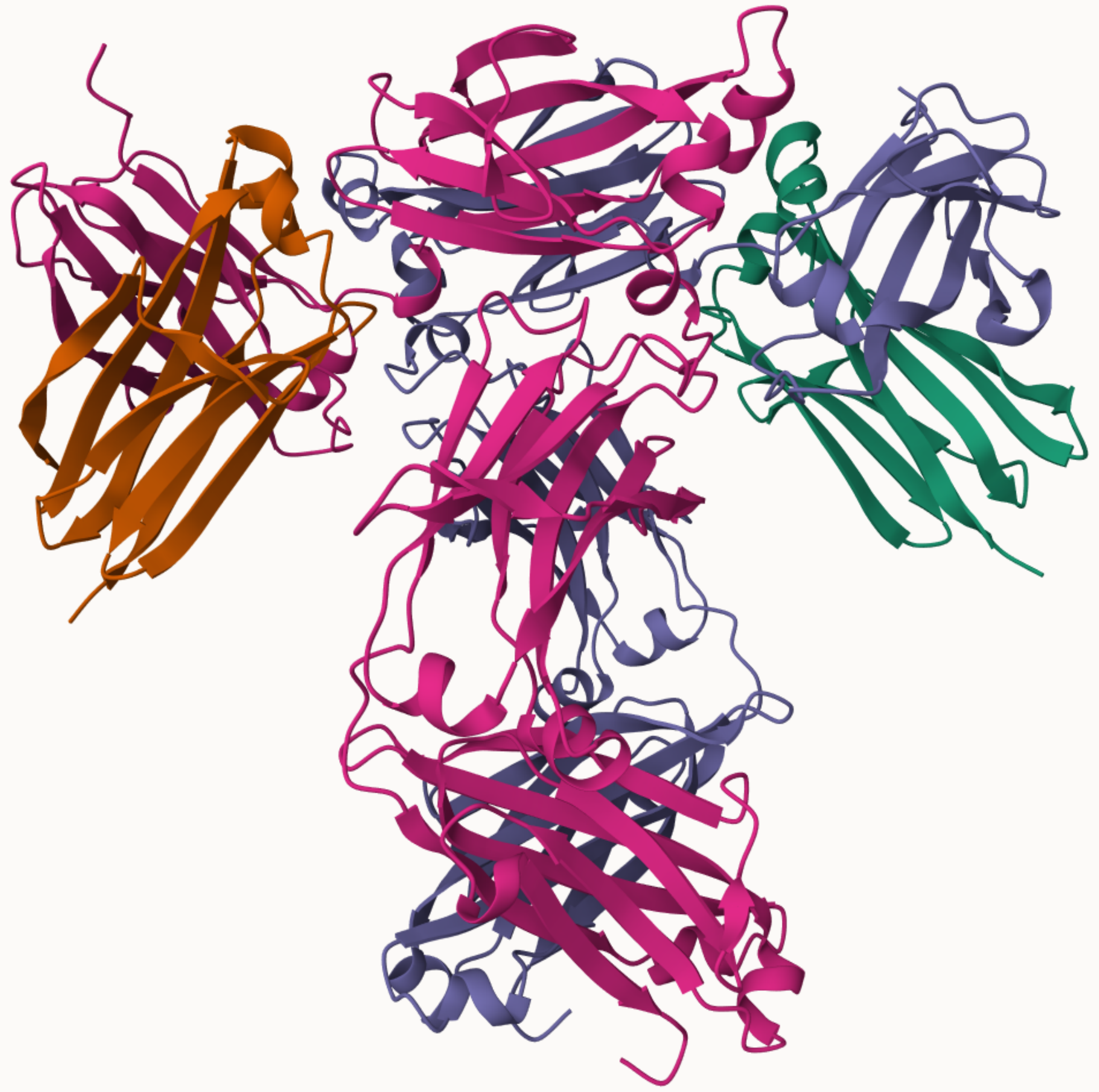

IgM4 Eublepharis macularius

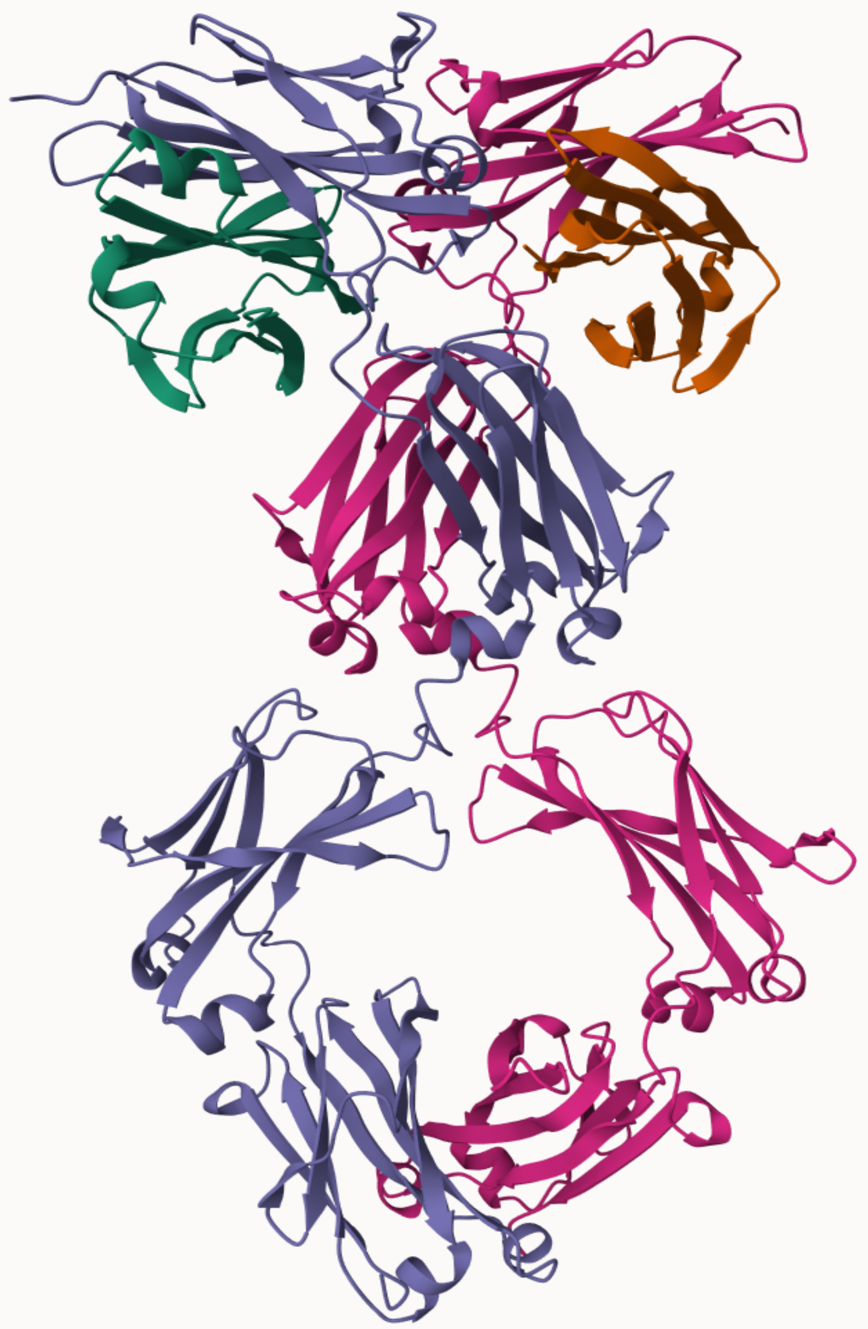

IgM5 Eublepharis macularius

### figures_03_conservation_profile.png

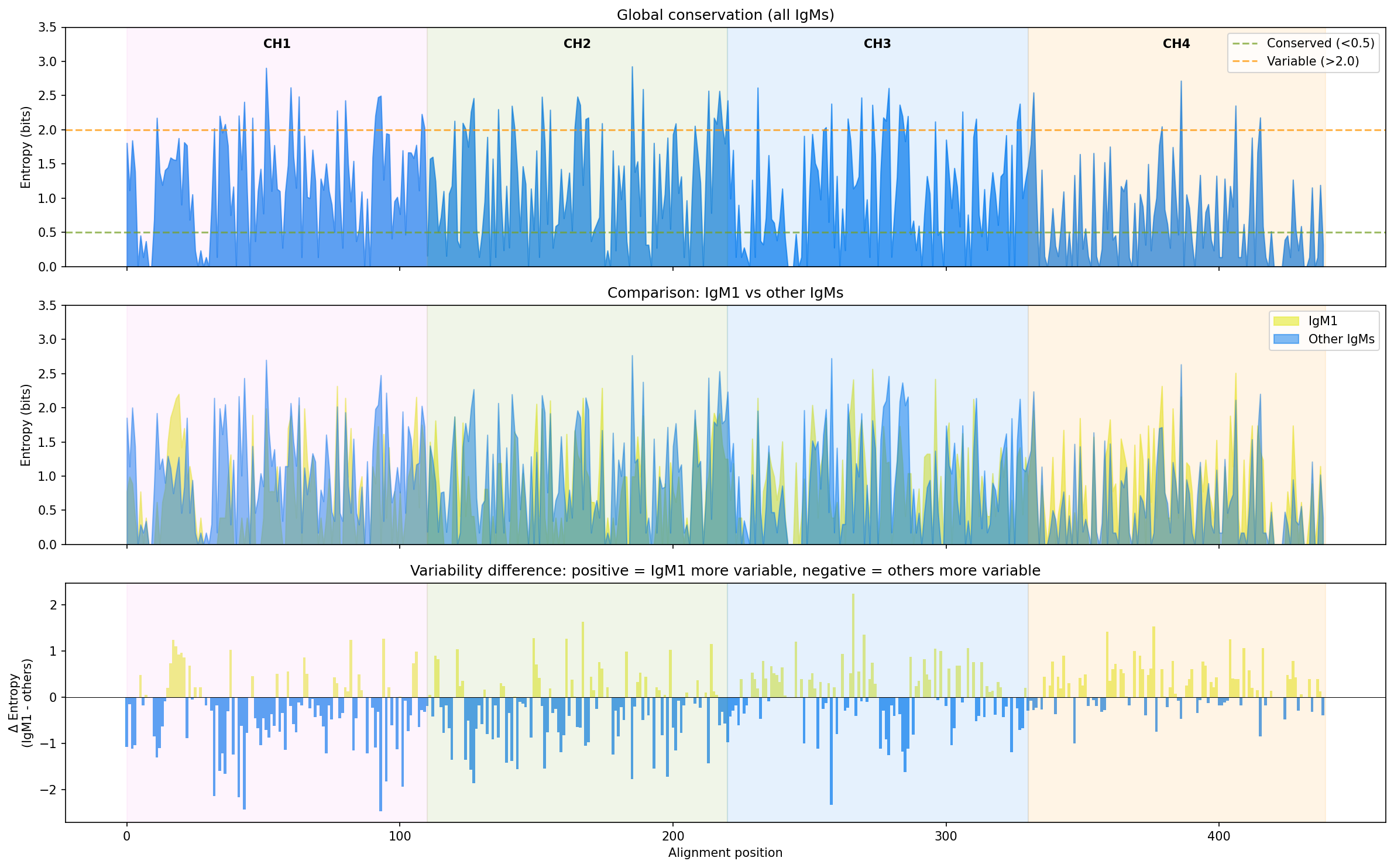

### figures_04_diagnostic_positions.png

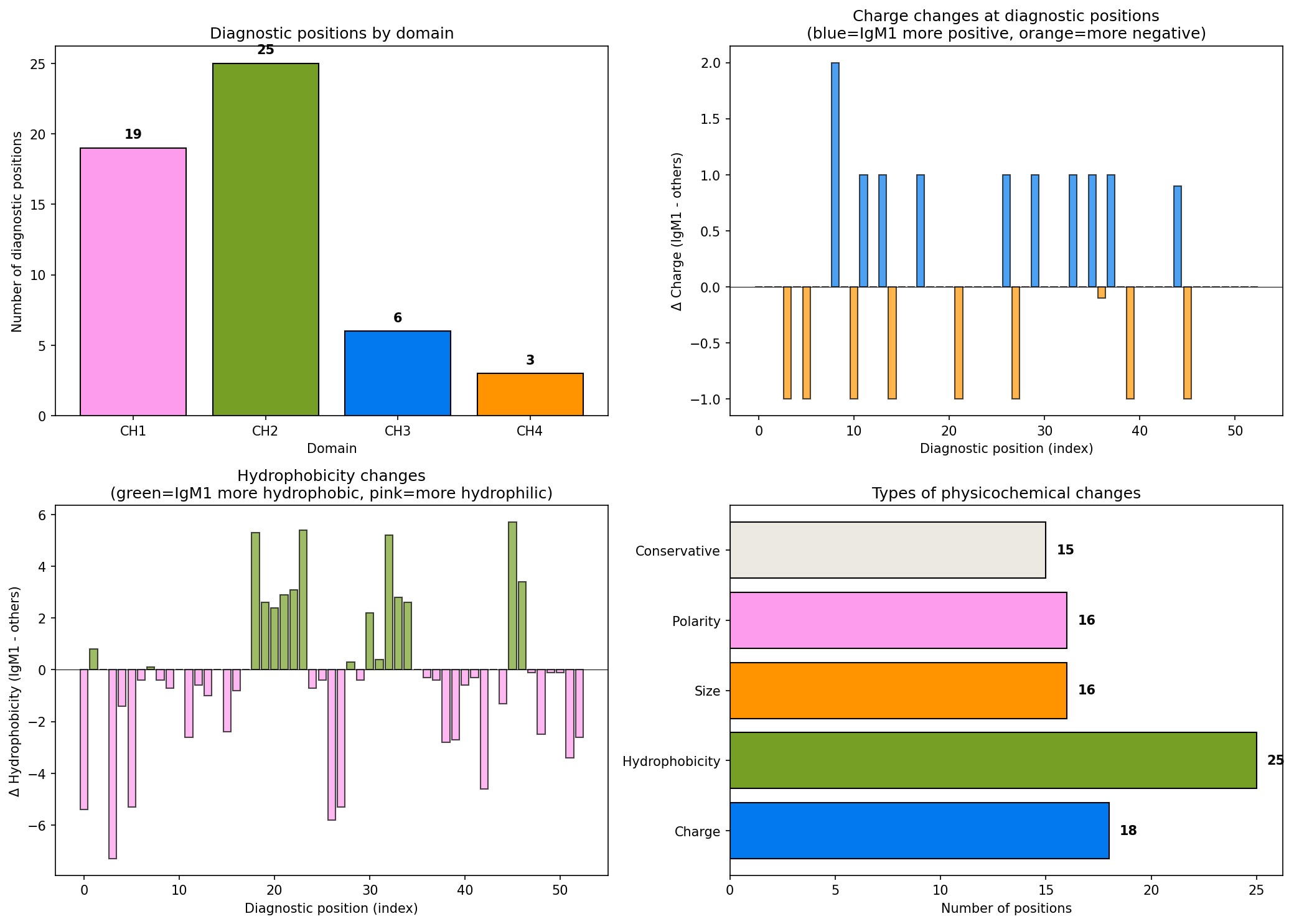

### figures_05_domain_properties.png

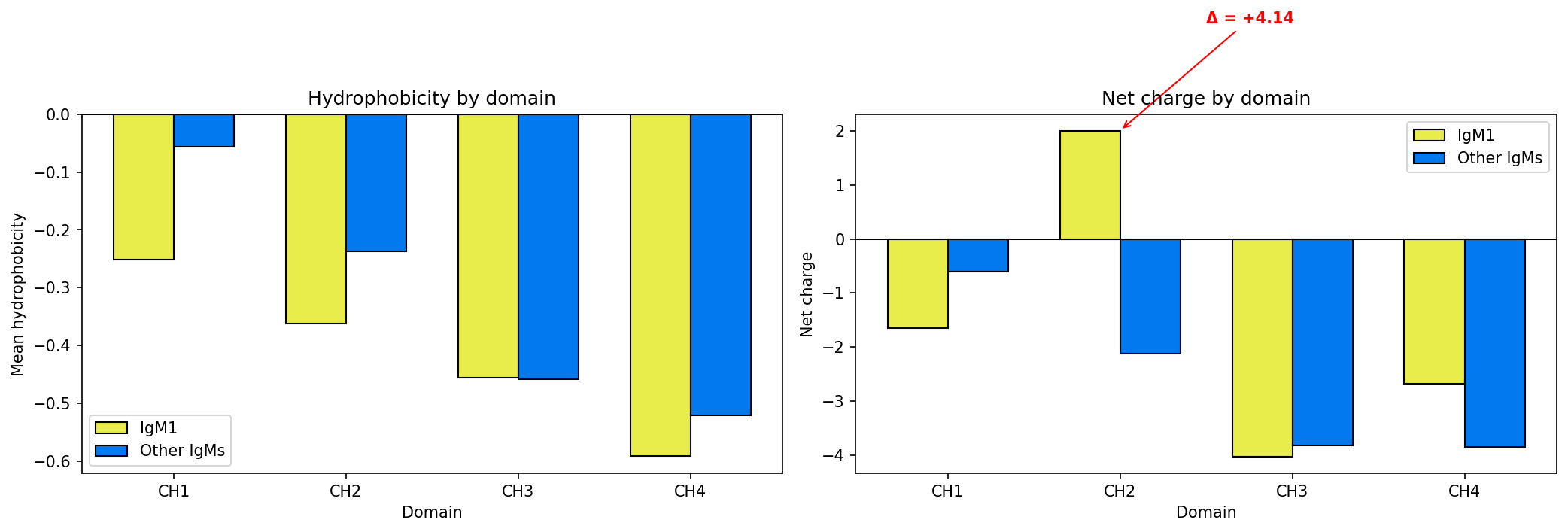

### figures_08_structural_analysis.pdf

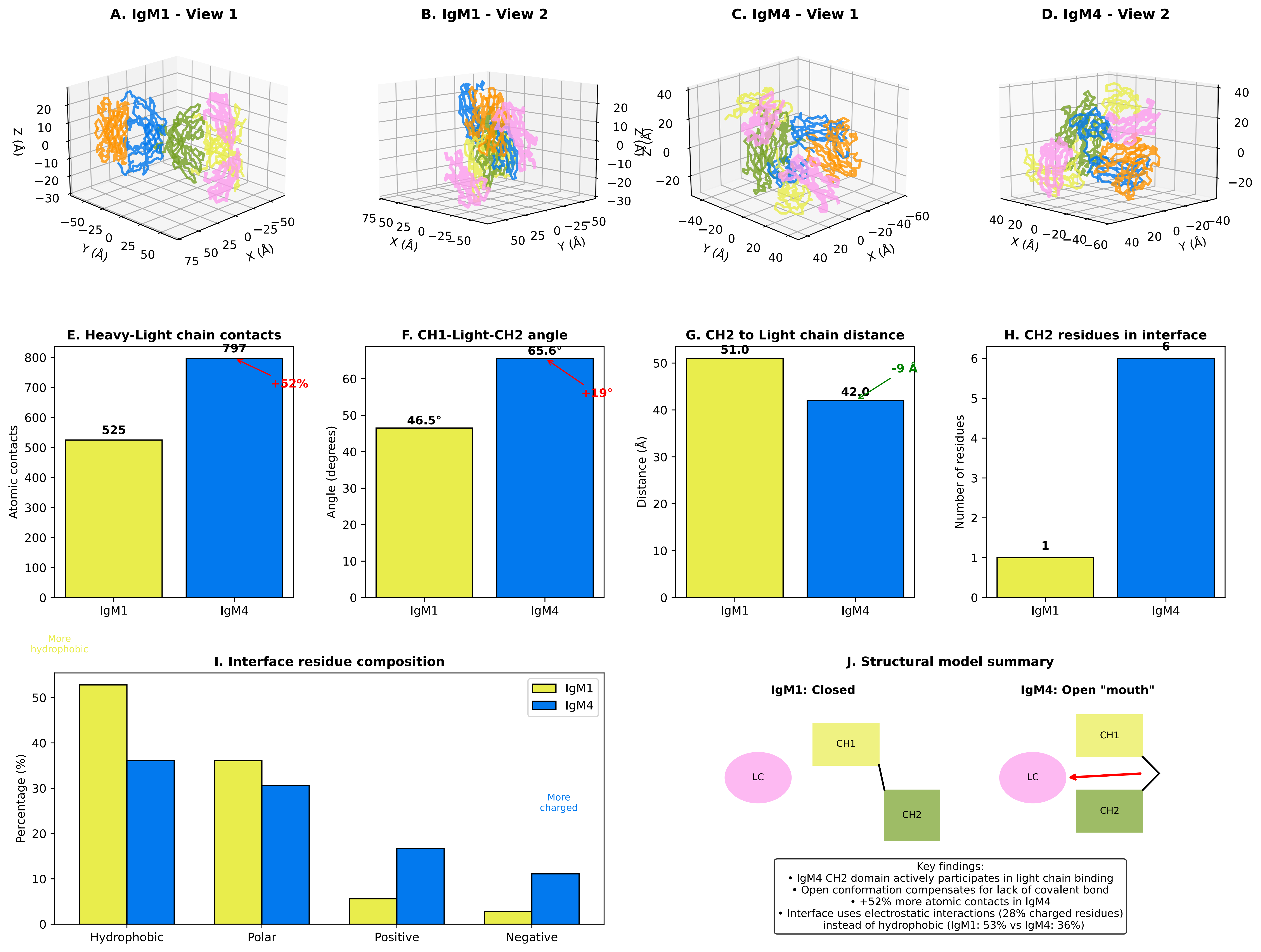

### tree.pdf

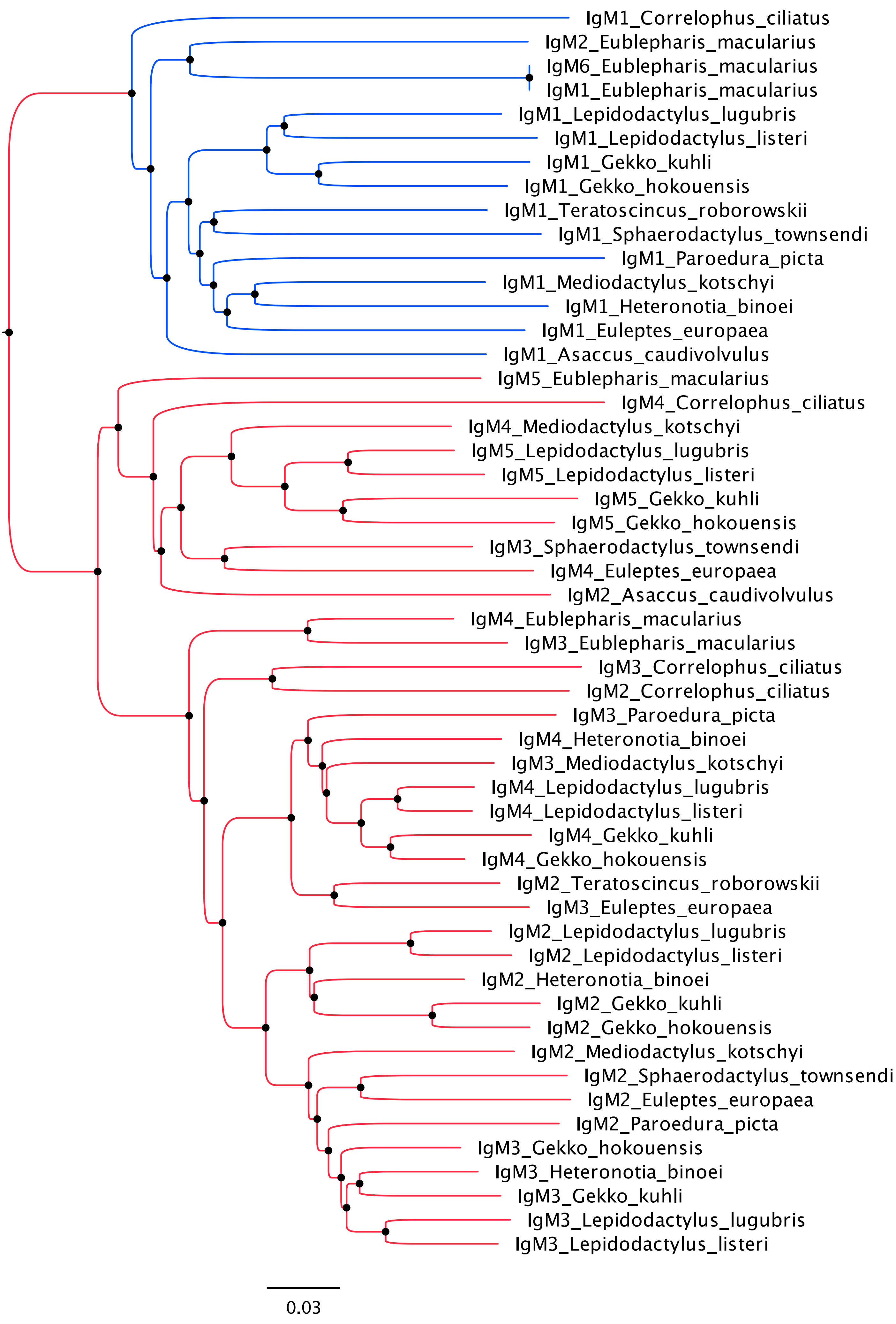
